# The MKK3 MAPK cascade integrates temperature and after-ripening signals to modulate seed germination

**DOI:** 10.1101/2024.01.28.577514

**Authors:** Masahiko Otani, Ryo Tojo, Sarah Regnard, Lipeng Zheng, Takumi Hoshi, Suzuha Ohmori, Natsuki Tachibana, Tomohiro Sano, Shizuka Koshimizu, Kazuya Ichimura, Jean Colcombet, Naoto Kawakami

## Abstract

Temperature is a major environmental cue for seed germination. The permissive temperature range for germination is narrow in dormant seeds and expands during after-ripening. Quantitative trait loci analyses of pre-harvest sprouting in cereals have revealed that MKK3, a mitogen-activated protein kinase (MAPK) cascade protein, is a negative regulator of grain dormancy. Here we show that the MAPKKK19/20-MKK3-MPK1/2/7/14 cascade modulates germination temperature range in Arabidopsis seeds by elevating germinability of the seeds at sub- and supra-optimal temperatures. The expression of *MAPKKK19* and *MAPKKK20* is regulated by an unidentified temperature sensing and signaling mechanism the sensitivity of which is modulated during after-ripening of the seeds, and MPK7 is activated at the permissive temperature for germination regulated by expression levels of *MAPKKK19/20*. Activation of the MKK3 cascade represses abscisic acid (ABA) biosynthesis enzyme gene expression, and induces expression of ABA catabolic enzyme and gibberellic acid biosynthesis enzyme genes, resulting in expansion of the germinable temperature range. Our data demonstrate that the MKK3 cascade integrates temperature and after-ripening signals to germination processes including phytohormone metabolism.

## Introduction

Temperature and seed dormancy are two important factors controling seed germination. Temperature is a major environmental factor and seed dormancy is an adaptive trait that enables the seeds to germinate in optimal season for vegetative and reproductive growth. It has been shown that seed responsiveness to temperature is closely related to the dormancy level in soil-buried seeds of winter and summer annuals (Baskin and Baskin, 2014). Primary dormancy of freshly harvested seeds gradually decreases with after-ripening, due to an expansion in the range of permissive germination temperatures (Baskin and Baskin 2014). In winter annual species such as Arabidopsis (*Arabidopsis thaliana* L. Heynh.), the seeds are dispersed from mother plants in spring, but at that time they do not germinate under any temperature conditions. From spring to autumn, the maximal permissive temperature for germination rises gradually during after-ripening, but germination is still suppressed by because temperatures are still higher than the upper limit for germination. So, the seeds do not germinate until the autumn when the temperature falls below the upper limit (Baskin and Baskin 2014). Therefore, the temperature sensing and signaling mechanism is modulated during after-ripening, and this allows the seeds to germinate in the appropriate season for their growth.

Abscisic acid (ABA) and gibberellic acid (GA) are the main phytohormones that antagonistically regulate seed germination. Studies have shown that supra-optimal high temperatures inhibit the germination of imbibed Arabidopsis and lettuce seeds by inducing expression of the ABA biosynthesis enzyme gene *NCED*, and repressing expression of the GA biosynthesis enzyme gene *GA3ox,* which increases ABA levels and decreases GA levels (Gonai et al. 2004, Toh et al. 2008, Argyris et al. 2008).

Pre-harvest sprouting (PHS) of maturing seeds severely reduces yield and quality of grain crops such as rice, wheat and barley (Singh et al. 2021). PHS tolerance has been shown to be closely linked with seed dormancy and regulated by quantitative trait loci (QTL). Several genes have been identified from major QTLs in rice (Sugimoto et al., 2010), wheat (Nakamura et al. 2011; Barrero et al. 2015) and barley (Nakamura et al. 2011; Barrero et al. 2015; Sato et al. 2016). Mitogen-activated protein kinase (MAPK) cascades are a common mechanism for transducing external and internal signals to cellular responses in eukaryotes. The MAPK cascades consist of at least three protein kinases, MAPK kinase kinase (MAPKKK), MAPK kinase (MKK), and MAPK (MPK), and are activated by consecutive phosphorylation (Ichimura *et al*., 2002). In plants, it has been reported that MAPK cascades are involved in various cellular processes such as biotic/abiotic stress responses, phytohormone responses, embryo development and plant growth (Xu and Zhang, 2015). The Arabidopsis genome codes for 80 MAPKKKs, 10 MAPKKs and 20 MPKs, and specific members of the families are involved in the specific signaling pathways (Ichimura et al. 2002, Jonak et al. 2002). MAPKKK activity is thought to be regulated by phosphorylation, for example in the case of immunity (Bi et al. 2018). Nevertheless, transcriptional regulation of clade III *MAPKKK*s seems to be the determinant of MKK3 module activation, explaining the delayed activation kinetics of its downstream group C MPKs (Colcombet at al. 2016). Recently, PHS QTL analyses of wheat and barley identified MKK3 as a negative regulator of seed dormancy (Nakamura et al. 2016; Torada et al. 2016). MKK3 has been reported to have multiple-functions in stress responses in both plantlets and adult plants (Colcombet et al. 2016). In the current study, we identified MKK3 containing MAPK cascade components which are involved in temperature signalling, and revealed their regulation mechanism and role in germination temperature range regulation in both freshly harvested and after-ripened Arabidopsis seeds.

## Results

### Arabidopsis MKK3 regulates germination response to temperature in freshly harvested and after-ripened seeds

We first analyzed the function of MKK3 on Arabidopsis seed dormancy and germination by using loss-of-function mutant alleles, *mkk3-1* and *mkk3-2* (Supplementary Fig. 1a; Takahashi et al. 2007, Sözen et al. 2020). Freshly harvested seeds of *mkk3-1* and *mkk3-2* showed slower speeds and lower percentages of germination than wild type (WT; Col-0) when imbibed at 22 °C (Supplementary Fig. 1b, c). The seeds of *mkk3-1* had a prolonged after-ripening period for coming out from dormancy (Fig. 1a), and their germination was stimulated by cold stratification (Fig. 1b). These observations indicate that *MKK3* works as a negative regulator of primary dormancy in Arabidopsis, as has been reported in wheat and barley (Torada et al. 2016, Nakamura et al. 2016).

**Fig. 1.**
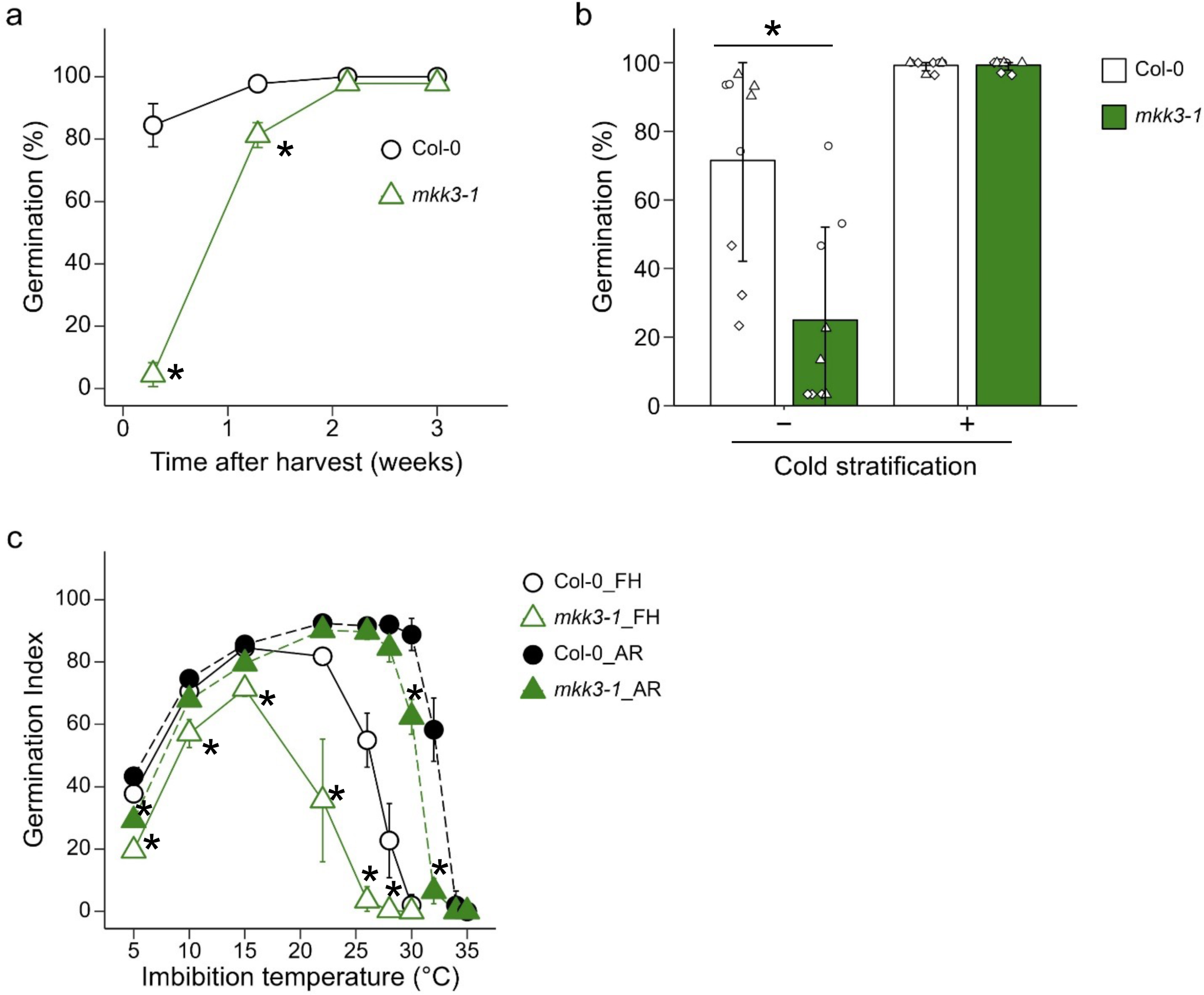
MKK3 is a modulator of germination response to temperature in both freshly harvested and after-ripened seeds. Asterisks indicate statistical differences between wild type (Col-0) and *mkk3-1* (p < 0.05, Student’s t test). **a** Enhanced dormancy phenotype of *mkk3-1* seeds. Freshly harvested seeds were stored in a desiccator at room temperature for up to 3 weeks. Seeds were imbibed at 22 °C for 7 days without cold stratification. Values are means of three technical replicates with SDs. We obtained similar results from triplicate experiments, and typical data are presented. **b** Effect of cold stratification on germination of the freshly harvested seeds. The seeds at DAH 2 were imbibed at 22 °C for 5 days with (+) or without (−) cold stratification at 4 °C for 4 days. Results from three independent seed batches are shown with averages and SDs. **c** MKK3 is a modulator of germination temperature range. Germination index of freshly harvested (FH, DAH 2, open symbols) and after-ripened (AR, DAH 150, closed symbols) seeds imbibed at 5 to 35 °C for 14 days are shown. Results from three independent seed batches are shown with averages and SDs.

We next analyzed germination at various temperatures by using freshly harvested (FH) and after-ripened (AR) seeds. The maximum germination ability, represented by germination index (GI), in FH seeds of WT was 15 °C, whereas in AR seeds the optimal germination temperature increased to around 26 °C, as commonly observed in winter-annual species (Fig. 1c, Baskin & Baskin 1983). The seeds of *mkk3-1* showed higher sensitivity to supra-optimal temperatures, with germination of FH and AR seeds respectively requiring ca. 6 °C and 2 °C lower temperatures, than WT. Also, at suboptimal temperatures, the germination speed of *mkk3-1* FH and AR seeds was clearly slower than WT (Fig. 1c for GI, Supplemental Fig. 1d, e for germination time course). These results suggest that MKK3 is a positive regulator of germination at both sub- and supra-optimal temperatures, and enables seeds to germinate over a range of temperatures.

### Expression of *MAPKKK19* and *MAPKKK20* is regulated by temperature during seed imbibition

It has been reported that transcriptional up-regulation of clade-III *MEKK-like MAPKKK*s by ABA is responsible for the activation of downstream kinases (Matsuoka et al. 2015, Danquah et al. 2015, Colcombet et al. 2016). In the same way, MKK3-MPK2 activation by wounding has been shown to depend on the transcriptional up-regulation of several clade-III *MAPKKK*s (Sözen 2020). In the current study, our transcriptome analysis (GSE229182) revealed that of all the clade-III *MAPKKK*s, only *MAPKKK19* and *MAPKKK20* were induced in germinating seeds (Supplementary Fig. 2). Expression of *MAPKKK19* was relatively high in dry seeds, but decreased to low levels after imbibition at supra-optimal temperatures in both FH (26 °C) and AR (34 °C) seeds (Fig. 2a and b). In germinating AR seeds imbibed at 26 °C, the expression levels were initially reduced during the first 6 h, but increased to high levels after 24 h. Expression of *MAPKKK20* was repressed in freshly harvested seeds at 26 °C, but temporarily induced in after-ripened seeds and peaked at 3h after the start of imbibition irrespective of temperature and germination (Fig. 2a and b). Also, *MAPKKK20* expression was re-induced in germinating AR seeds imbibed at the optimal temperature, peaking at 24 h after the start of imbibition, but was repressed in non-germinating FH and AR seeds imbibed at supra-optimal temperatures.

**Fig. 2.**
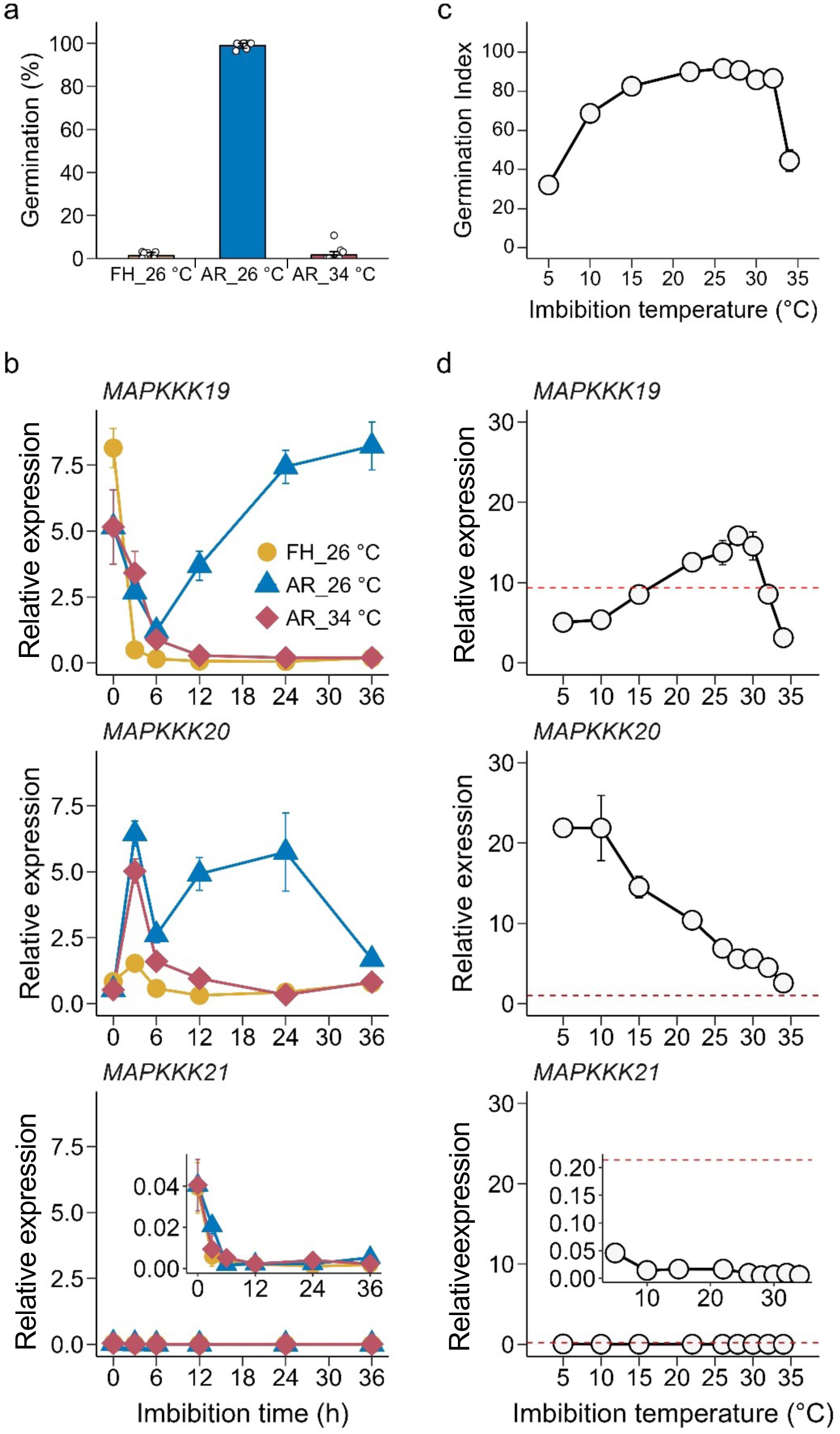
Temperature dependent expression *MAPKKK19* and *MAPKKK20* in FH and AR seeds. **a** Final percentage germination of FH (DAH 2) and AR (DAH 49) Col-0 seeds used for gene expression analysis in **b**. The FH seeds were imbibed at 26 °C, and the AR seeds were imbibed at either 26 °C or 34 °C for 7 days. Results from three technical replicates are shown with averages and SDs. **b** Effect of imbibition temperature on the expression time course of *MAPKKK19, 20* and *21* in FH and AR seeds. FH seeds imbibed at 26 °C (orange circles), and AR seeds imbibed at 26 °C (blue triangles) or 34 °C (red diamonds) were used for RNA extraction. **c** GI of AR (DAH 330) Col-0 seeds used for gene expression analysis in **d**. The seeds were imbibed at various temperatures for 14 days. Values are means of three technical replicates with SDs. **d** Effect of imbibition temperature on the expression of *MAPKKK19, 20* and *21*. Total RNA was prepared from 24 h imbibed seeds. Dashed red lines indicate the expression levels in dry seeds. **b** and **d** Transcript levels were quantified by qRT-PCR. We obtained similar results from three biological replicates, and typical data is presented.

In concert with germination ability (GI), the maximum expression of *MAPKKK19* in 24 h imbibed AR seeds was at around 26 °C and reduced at supra- and sub-optimal temperatures (Fig. 2c and d). Furthermore, the expression of *MAPKKK20* was also temperature-dependent, with maximum expression at 5 °C but steadily decreasing as the temperature rose (Fig. 2d).

### *MAPKKK19* and *MAPKKK20* are involved in the regulation of seed germination in response to temperature

In order to understand the role of *MAPKKK19* and *MAPKKK20* on seed germination response to temperature, we isolated DNA insertion mutants. We also isolated DNA insertion mutant of *MAPKKK21* which is a closest paralog of *MAPKKK19* and *MAPKKK20* to elucidate the possibility of redundant function of the genes (Supplementary Fig. 2a). Expression of *MAPKKK21* was very low in the imbibed FH and AR seeds at any temperature conditions (Fig. 2b, d), but the expression was evident during seed development (Supplementary Fig. 4). *mapkkk19-1* and *mapkkk21-1* were transposon insertion lines of the Nossen ecotype, and were backcrossed with Col-0 (Supplementary Fig. 3). *mapkkk20-3* was a T-DNA insertion line of Col-0. The FH seeds of the single mutants showed almost the same germination as wild type at 22 °C, but the seeds of the double mutants showed lower germination percentage than WT (Fig. 3a). Furthermore, the FH seeds of *mapkkk19/20/21* triple mutant showed a lower germination percentage than WT and the double mutants (Fig. 3a). *mapkkk19/20/21* seeds had a prolonged after-ripening period, and their germination was stimulated by cold stratification as observed in *mkk3-1* seeds (Supplementary Fig. 5a and b). These results suggest that all the three *MAPKKK*s are involved in the regulation of seed dormancy.

**Fig. 3.**
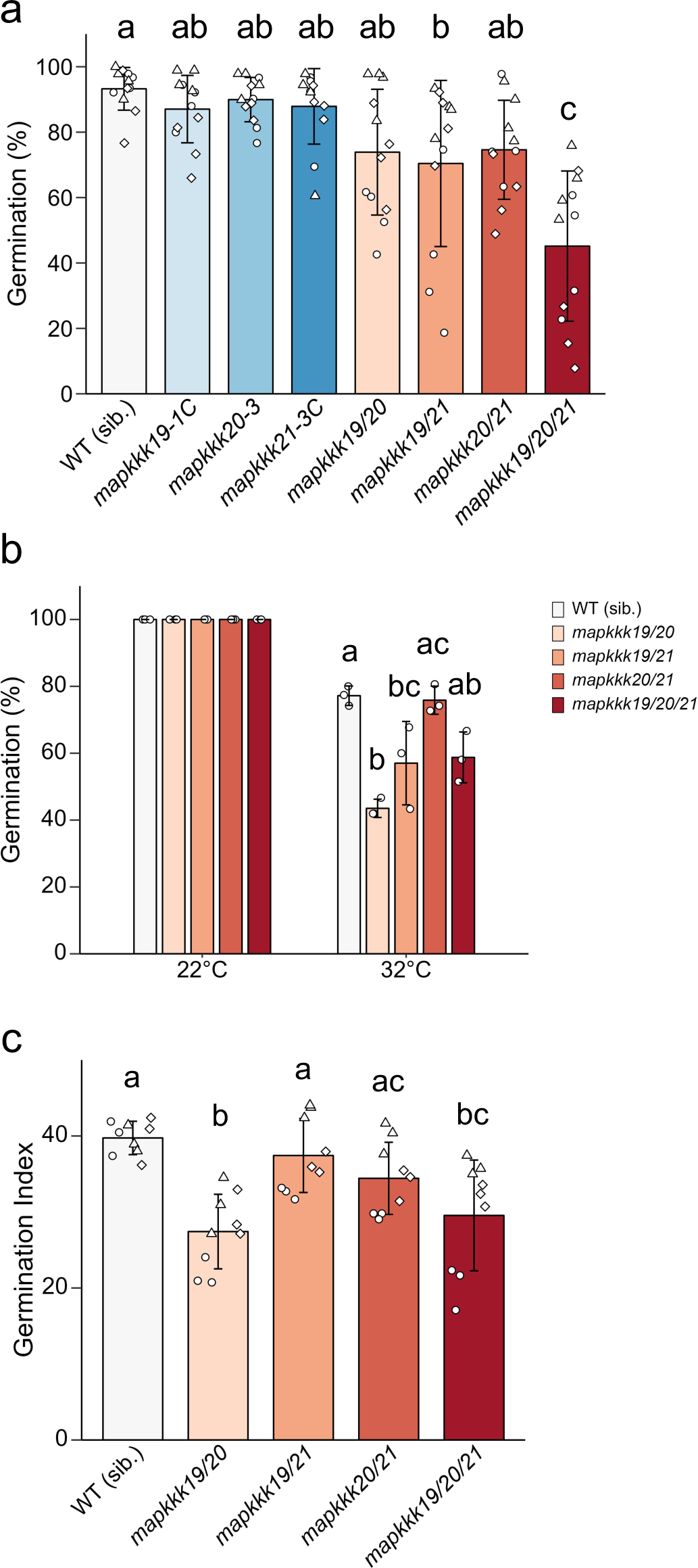
*MAPKKK19* and *20* are responsible for germination of FH and AR seeds, and *MAPKKK21* contributes to germination of FH seeds. Significant differences between samples are indicated by different letters (Tukey HSD tests, PL<L0.05). **a** Germination of FH seeds of single and multiple mutants at 22 °C. Freshly harvested (DAH 2) seeds were imbibed for 7 days. Results from three biological replicates are shown with averages and SDs. **b** Effect of supra-optimal temperature on germination of AR seeds. The seeds were imbibed at 22 °C and 32 °C for 5 days. We obtained similar results from three biological replicates, and a typical result is shown. **c** Effect of sub-optimal temperature on germination of AR seeds. The seeds were imbibed at 5 °C for 14 days. GI values were presented as mean (±SD) of three biological replicates.

We next analyzed the function of the MAPKKKs on germination response to temperature with AR seeds. At supra-optimal temperature (32 °C), the seeds of *mapkkk19/20* showed significantly lower germination percentage than WT (Fig. 3b). In contrast to the FH seeds, AR seeds of *mapkkk19/20/21* showed similar germination to *mapkkk19/20* (Fig. 3b). At sub-optimal temperature (5 °C), the germination speed of *mapkkk19/20* seeds was slower than WT, and the delayed germination phenotype was not enhanced in *mapkkk19/20/21* seeds (Fig. 3c, Supplementary Fig. 5c). These results suggest that *MAPKKK19* and *MAPKKK20* are responsible for the germination response to temperature in both FR and AR seeds, but *MAPKKK21* is no longer effective in AR seeds.

We also analyzed the contribution of other clade III MAPKKKs on germination of FH and AR seeds. MAPKKK13 and MAPKKK14 are known to have putative transmembrane motifs at C-terminal domains (Schwacke et al. 2003, Sözen et al 2020). The gain-of-function mutant, *mapkkk14-1*, produces a mutant protein that lacks a C-terminal transmembrane domain, and has been reported to show higher MPK2 activation ability than WT in response to wounding (Sözen et al. 2020). However, in our study the seeds of *mapkkk14-1* showed no germination phenotypes (Supplementary Fig. 6a to c). We also isolated the *mapkkk13-1* allele which has T-DNA insertion between kinase and the transmembrane domains, and produced a *mapkkk13-1/14-1* double mutant. However, the FH and AR seeds also showed very similar germination to WT (Supplementary Fig. 6c). We next used the gene editing loss-of-function alleles of *MAPKKK13* and *MAPKKK14* (Sözen et al. 2020; Regnard et al. 2024), but again the germination response to temperature of the FH and AR seeds of *map3k13CR/14CR* double mutant was very similar to WT (Supplementary Fig. 6d). Furthermore, the seeds of *mapkkk15/16* and *mapkkk17/18* also showed similar germination response to temperature as WT (Supplementary Fig. 7, 8). Therefore, these results suggest that *MAPKKK13* to *MAPKKK18* have almost no function in seed germination response to temperature.

To evaluate the contribution of transcriptional regulation of *MAPKKK*s on seed germination response to temperature, we analyzed *MAPKKK20* overexpression (*MAPKKK20^OX^*) lines (Supplementary Fig. 9a). The two independent lines accumulated ca. 36-fold more *MAPKKK20* transcripts in dry seeds than the non-transformant wild type (Supplementary Fig. 9b). The FH and AR seeds of *MAPKKK20^OX^* lines showed significantly higher percentage germination than WT at supra-optimal temperatures (Fig. 4a and b). These results suggest that transcriptional regulation of *MAPKKK20* plays an important role in germination response to temperature. On the other hand, *MAPKKK20* overexpression had almost no effect on germination at 5 oC (Supplementary Fig. 9c). The low temperature induced expression nature of the native *MAPKKK20* may mask the effect of the trans gene (Fig. 2d).

**Fig. 4.**
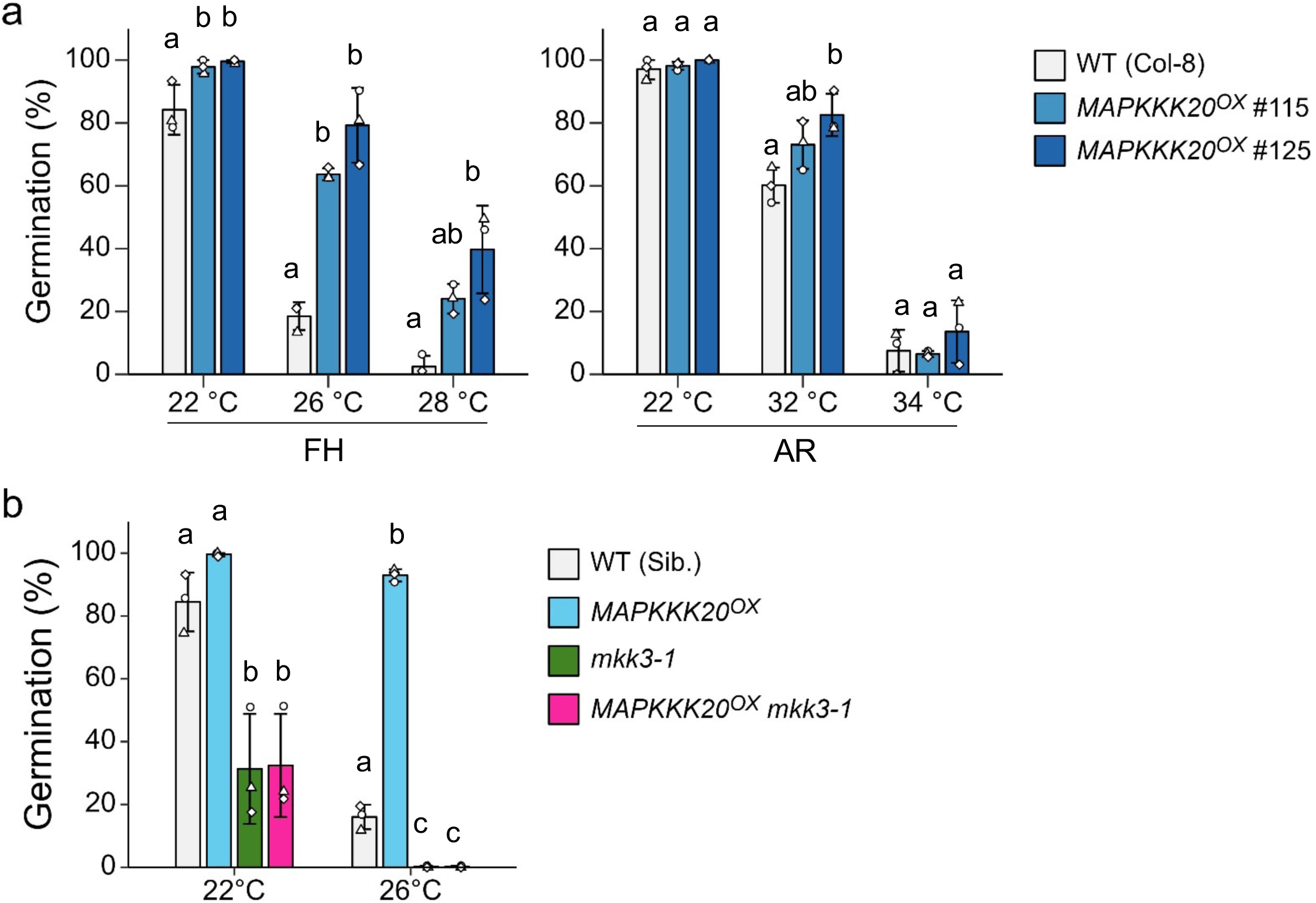
*MAPKKK20* over-expression stimulates germination of FH and AR seeds at supra-optimal temperatures in the MKK3 pathway. The values are the means (±SD) of three biological replicates. Significant differences between lines are indicated by different letters (Tukey HSD tests, PL<L0.05). **a** FH (DAH 2) and AR (DAH 64) *35S::MAPKKK20-10xMYC* (*MAPKKK20^OX^*) seed germination response to imbibition temperature. The seeds were imbibed for 5 days. **b** *mkk3-1* is epistatic to *MAPKKK20^OX^*. Freshly harvested (DAH 2) seeds were imbibed at 22 and 26 °C for 7 days without stratification.

We analyzed the genetic interaction between *MAPKKK20* and *MKK3* by creating a *MAPKKK20^OX^ mkk3-1* double mutant. In contrast to *MAPKKK20^OX^* seeds, the double mutant seeds showed lower percentage germination than WT, having almost the same germination rate as *mkk3-1* (Fig. 4b). This epistatic nature of *mkk3-1* suggests that MAPKKK20 works upstream of MKK3 for germination.

### Group C MPKs are involved in the regulation of seed germination response to temperature

It has been reported that MKK3 activates group C MPKs (i.e. MPK1, MPK2, MPK7 and MPK14) by direct binding (Dóczi et al. 2007, Lee et al. 2008, Matsuoka et al. 2015, Danquah et al. 2015). To identify MPKs which are involved in the regulation of seed germination response to temperature, we isolated and analyzed multiple knockout mutants of the group C *MPK*s (Supplementary Fig. 10). Among the four single mutants, only *mpk7-1* FH seeds showed lower percentage germination than WT at 22 °C (Fig. 5a). The seeds of *mpk1/2/14* triple mutant showed almost the same germination as WT, but multiple mutant seeds containing *mpk7-1* showed reduced germination (Fig. 5a). Among these multiple mutants, the seeds of *mpk1/2/7, mpk1/7/14* and *mpk1/2/7/14* showed greatly reduced germination phenotype (Fig. 5a). Dormancy of the *mpk1/7/14* and *mpk1/2/7/14* seeds was alleviated by after-ripening and cold-stratification treatment, similar to *mkk3* (Supplementary Fig. 11). These results indicate that *MPK7* has a prominent role, but that other group C MPKs redundantly work on germination of FH seeds, i.e. dormancy.

**Fig. 5.**
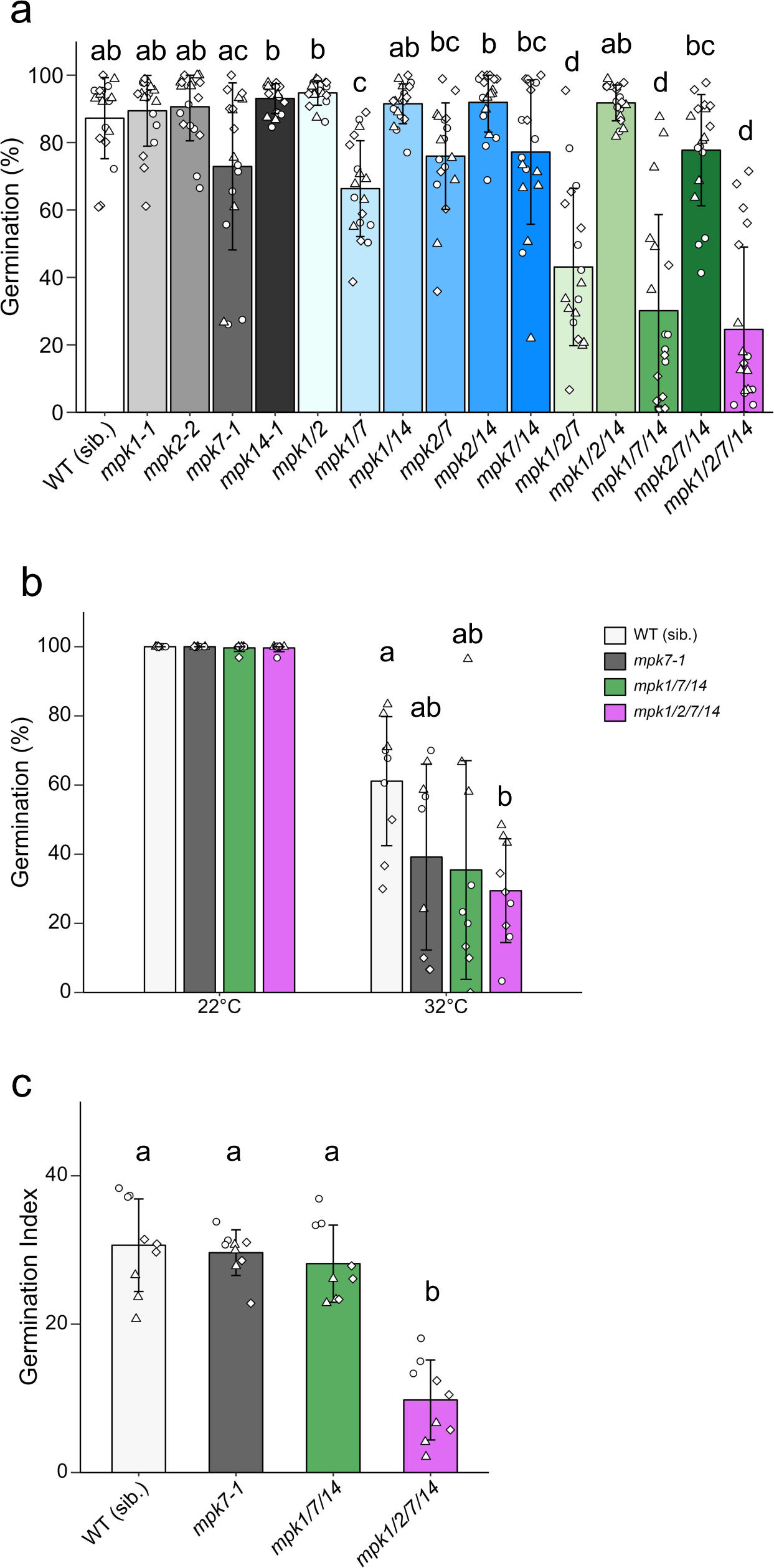
Group C MPKs are responsible for germination of FH and AR seeds. The values are the means (±SD) of three biological replicates. Significant differences between lines are indicated by different letters (Tukey HSD tests, PL<L0.05). **a** Germination of FH (DAH 2) seeds of single and multiple mutants at 22 °C. The seeds were imbibed for 7 days without cold stratification. **b** Effect of supra-optimal temperature on germination of AR seeds. The seeds were imbibed at either 22 °C or 32 °C for 5 days. **c** Effect of sub-optimal temperature on germination of AR seeds. Germination index of after-ripened seeds imbibed at 5 °C for 14 days.

The AR seeds of *mpk1/7/14* and *mpk1/2/7/14* were more sensitive to supra-optimal temperature than WT (Fig. 5b). At sub-optimal temperature (5 °C), the seeds of *mpk7-1* and *mpk1/7/14* showed a 1-day delay in germination, and the *mpk1/2/7/14* seeds showed a 3-day delay when compared with WT (Fig. 5c, Supplementary Fig. 11c). These results indicate that MPK7 and other group C MPKs redundantly work on promoting germination at supra-optimal temperatures. At sub-optimal low temperatures, MPK2 may have a major role, while other group C MPKs have redundant function on germination.

### MPK7 activity is regulated by MAPKKK19/20-MKK3 module in response to temperature

To understand the relationship between *MAPKKK19/20* expression, MPK activation and seed germination, we first analyzed MPK7 activity in germinating and non-germinating seeds since MPK7 was shown to have a major role in the regulation of germination (Fig. 5a, b). Germinating WT seeds showed a peak in MPK7 activity after 12 to 24 h of imbibition, but this increased activity was not observed in non-germinating FH and AR seeds imbibed at supra-optimal temperatures (Fig. 6a, Supplementary Fig. 12a). A peak in MPK7 activity was also observed in after-ripened seeds imbibed for 3h, irrespective of the imbibition temperature, but not detected in dormant seeds. This MPK7 activity was synchronized with the expression of *MAPKKK20* (Fig. 2b). However, this activation at 3 h was not detected in *mapkkk19/20* and *mapkkk19/20/21* seeds (Fig. 6b and c, Supplementary Fig. 12b and c). Therefore, the early activation of MPK7 appears to be induced by the expression of *MAPKKK20*, but it may not be enough for the completion of germination. These results suggest that activation of group C MPK after 12 to 24 h of imbibition is responsible for the germination response to temperature. We detected almost no MPK7 activity in germinating *mkk3-1* and *mapkkk19/20* seeds throughout the imbibition period (Fig. 6b). Unexpectedly, some MPK7 activity was detected after 1 h of imbibition in the *mapkkk19/20/21* seeds, but the activity diminished during subsequent imbibition (Fig. 6b and c), suggesting that the recorded activity might be the result of the complementation effect of other clade III *MAPKKK* genes expressed during development of the triple mutant seeds. These results suggest that MKK3, MAPKKK19 and MAPKKK20 are responsible for the activation of group C MPKs during imbibition, but MAPKKK21 is not. The contribution of MAPKKK21 was clearly observed for germination of freshly harvested seeds but not for germination of after-ripened seeds (Fig. 3), suggesting that MAPKKK21 may be activated only during seed development, and it work on germination of freshly matured seeds.

**Fig. 6.**
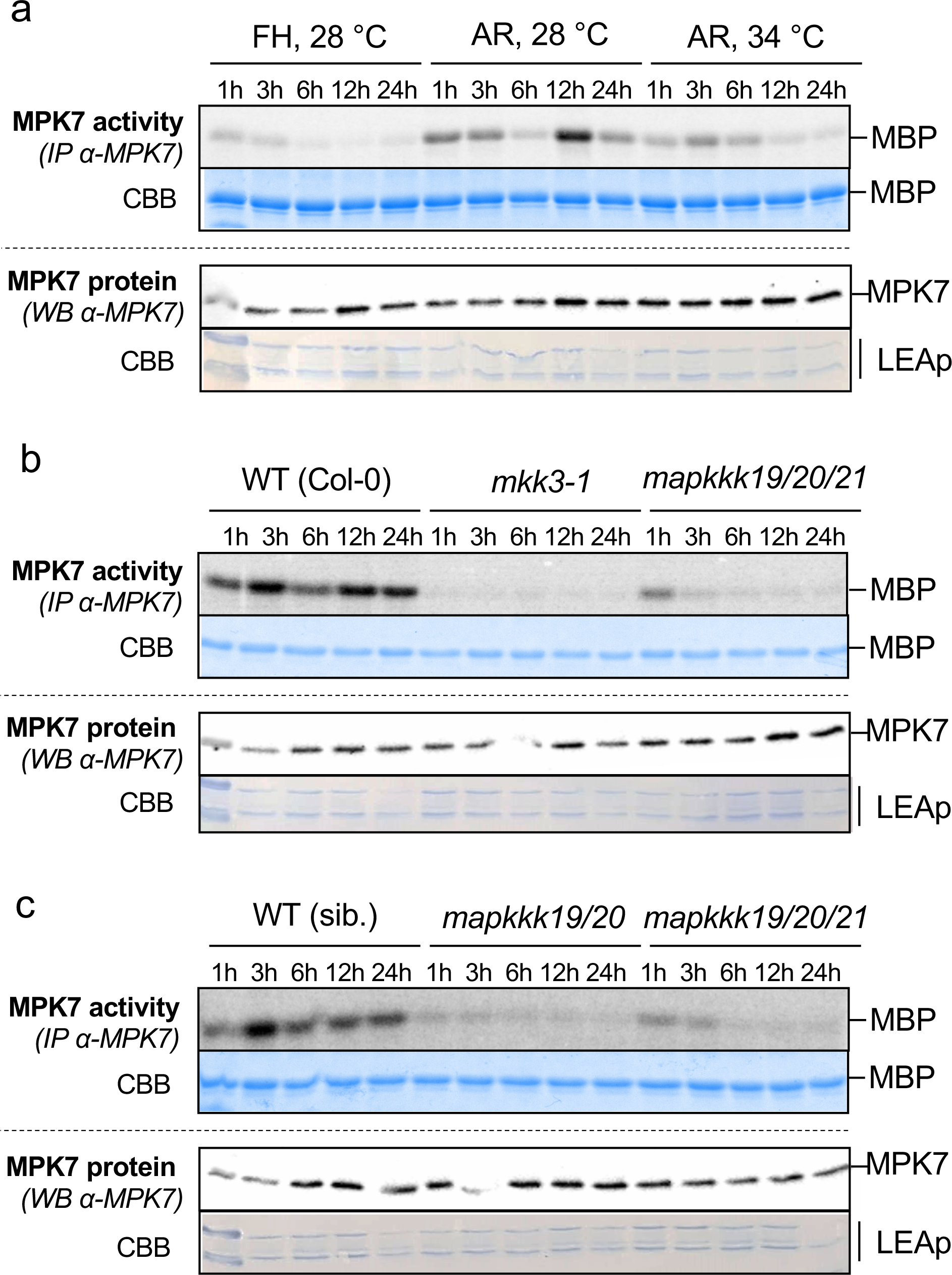
MPK7 activity in germinating and non-germinating seeds. MBP was used as a phosphorylation substrate of MPK7 which was immunoprecipitated from imbibed seeds with an anti-MPK7 antibody. Amount of MPK7 protein was monitored by immunoblot using an anti-MPK7 antibody. Equal loading was indicated by Coomassie staining of the LEA protein (LEAp) on the membrane. **A** Effect of imbibition temperature and seed dormancy status (after-ripening) on MPK7 activity. **b** MPK7 activity in after-ripened *mkk3-1* and *mapkkk19/20/21* seeds imbibed at permissive temperature for germination (28 °C). **c** MPK7 activity in after-ripened *mapkkk19/20* and *mapkkk19/20/21* seeds imbibed at permissive temperature for germination (28 °C).

### Effect of ABA and GA in *MAPKKK19/20* expression

It has been reported that high temperature inhibits seed germination by inducing ABA biosynthesis and suppressing GA biosynthesis in Arabidopsis and lettuce seeds (Toh et al. 2008, Argyris et al. 2008). So, we analyzed the effect of endogenous and exogenously applied ABA and GA on the expression of *MAPKKK19/20*. FH seeds of the ABA deficient *aba2-2*, showed almost no dormancy and germinated well at 28 °C, but this germination was inhibited by the application of ABA (Fig. 7a). Expression of *MAPKKK19* and *MAPKKK20* was repressed in the imbibed WT dormant seeds, but was de-repressed in *aba2-2* seeds (Fig. 7b). The expression of *MAPKKK19* and *MAPKKK20* in *aba2-2* seeds was moderately suppressed by the exogenously applied ABA. However, the expression of both *MAPKKK19* and *MAPKKK20* was not apparently affected in the seeds treated with the ABA biosynthesis inhibitor, fluridone, and ABA (Fig. 7c and d). These results suggest that the expression of *MAPKKK19* and *MAPKKK20* is not directly regulated by ABA, but the expression is controlled by temperature and physiological status of the seeds.

**Fig. 7.**
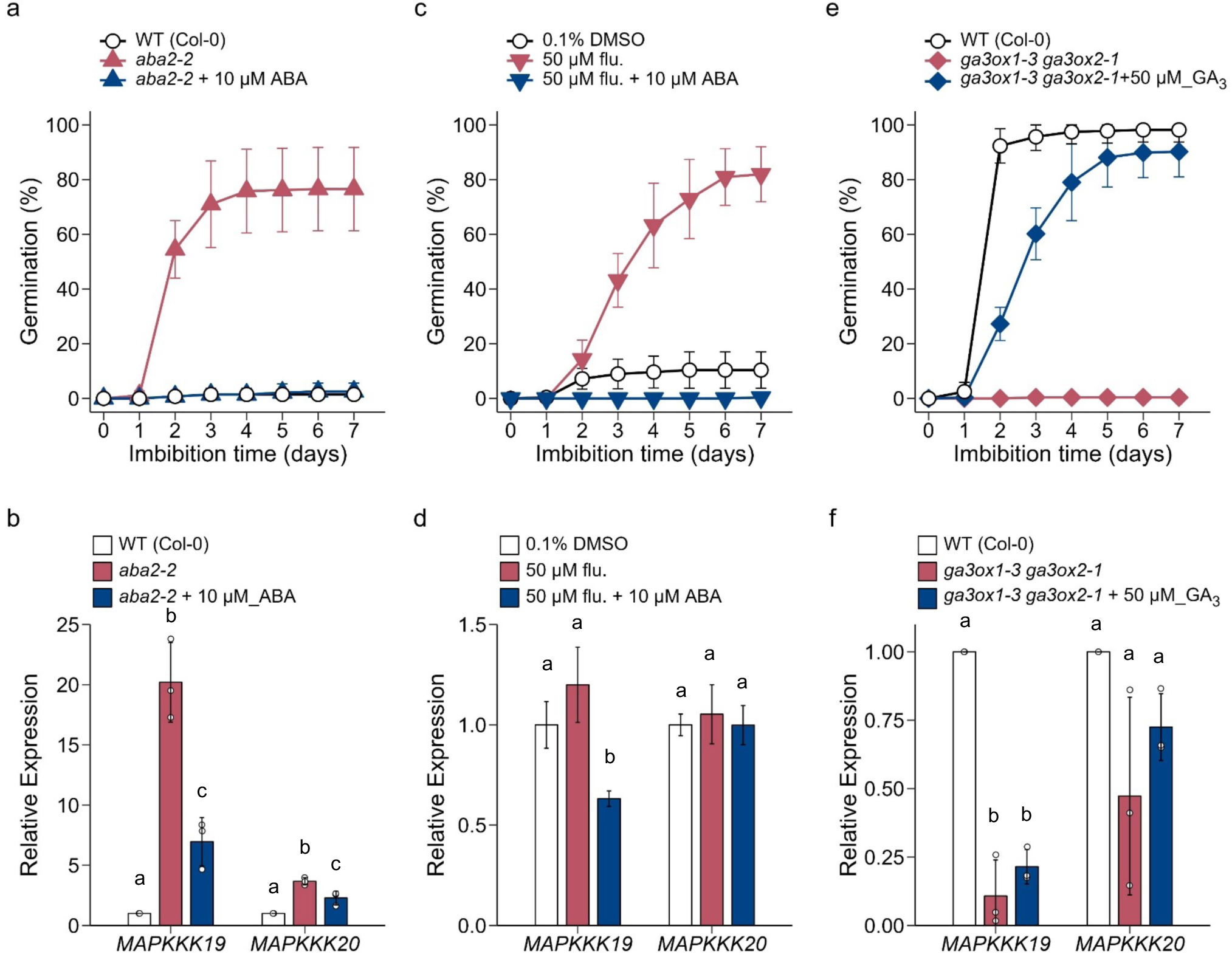
Effect of ABA and GA on the expression of *MAPKKK19* and *MAPKKK20*. Seed germination and gene expression data are the means (±SD) of three biological replicates. Total RNA was prepared from FH (**a**; DAH2) or AR (**c**; DAH 80, **e**; DAH 50) seeds imbibed for 24 h or 48 h. Transcript levels were quantified by qRT-PCR with At2g20000 as an internal standard. Significant differences in the gene expression levels are indicated by different letters (Tukey HSD tests, PL<L0.05). **a**-**d** Effect of endogenous and exogenous ABA on seed germination (**a, c**) and gene expression (**b, d**). The endogenous ABA levels in FH seeds imbibed at 28 °C were reduced by *aba2-2* mutation (**a, b**) or by the ABA biosynthesis inhibitor, fluridon in Col-0 AR seeds imbibed at 32 °C (**c** and **d**). Total RNA was prepared from seeds imbibed for 24 h. **e, f** Effect of endogenous and exogenous GA on seed germination (**e**) and gene expression (**f**). AR seeds were imbibed at 30 °C. Total RNA was prepared from AR seeds imbibed for 24h.

After-ripened GA deficient, *ga3ox1-3 ga3ox2-1* double mutant seeds imbibed at 30 °C could not germinate like similarly imbibed WT seeds, but application of exogenous GA_3_ enabled the double mutant seeds to germinate (Fig. 7e). Expression of *MAPKKK19* was repressed in the GA deficient mutant seeds, and this repression was not reversed by the application of exogenous GA (Fig. 7f). In addition, we could not detect any significant effect of the GA deficiency or exogenous GA application on the expression of *MAPKKK20* (Fig. 7f). These results suggest that the expression of *MAPKKK19* and *MAPKKK20* is not regulated by GA.

### MKK3-MAPK cascade modulates ABA and GA metabolism

To understand the molecular mechanism of germination regulation by the MKK3-MAPK cascade, we analyzed the expression of ABA and GA metabolism enzyme genes in the seeds of *MAPKKK20* over-expression lines. In this experiment, *MAPKKK20^OX^* seeds showed higher percentage germination than WT at 34 °C (Fig. 8a). It has been reported that the key ABA biosynthesis enzyme genes, *NCED2, NCED5* and *NCED9* are induced under supra-optimal temperature conditions (Toh et al. 2008, Fig. 8b). The expression of all the three *NCED*s was reduced in *MAPKKK20^OX^* seeds imbibed at supra-optimal temperature, 34 °C, when compared with WT (Fig. 8b). The ABA catabolism enzyme genes, *CYP707A1*, *CYP707A2* and *CYP707A3*, have been reported to be involved in germination, and *CYP707A2* has a major role in the rapid decrease in ABA content right after imbibition (Kushiro et al. 2004, Okamoto et al. 2006). At the permissive 26 °C temperature, all three ABA catabolism enzyme genes showed significantly higher expression levels in *MAPKKK20^OX^* seeds than in WT (Fig. 8c). Expression levels of *CYP707A2* and *CYP707A3* were also up-regulated at supra-optimal 34 °C temperature in *MAPKKK20^OX^*seeds (Fig. 8c). These results suggest that the MKK3-MAPK module stimulates seed germination by reducing ABA levels through repression of ABA biosynthesis genes and inducing ABA catabolism genes.

**Fig. 8.**
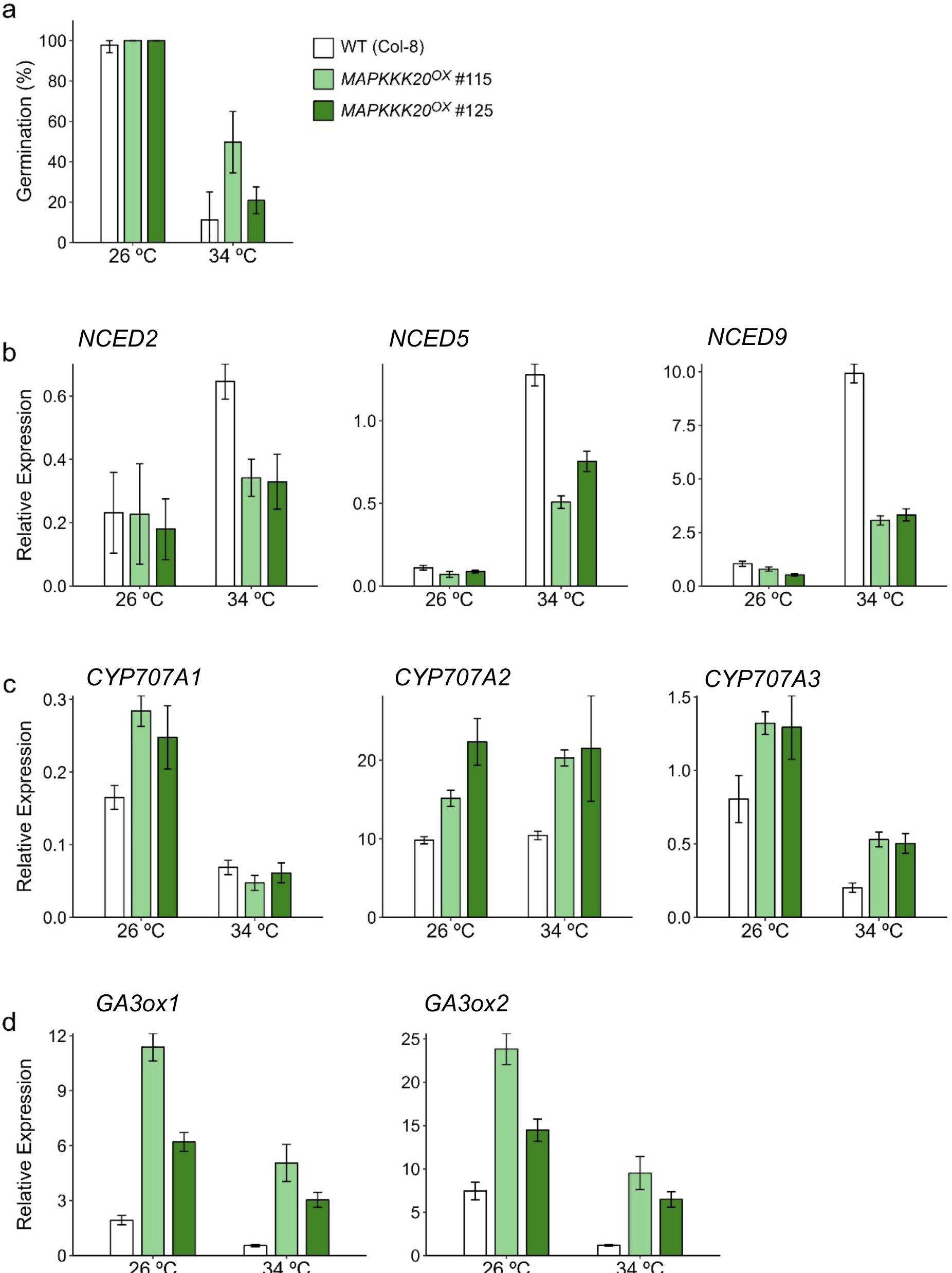
*MAPKKK20* can induce expression of ABA catabolism and GA biosynthesis genes and reduce expression of ABA biosynthesis genes. Values are means of three technical replicates with SDs. We obtained similar results from triplicate experiments, and typical data are presented. Transcript levels were quantified by qRT-PCR with At2g20000 as an internal control for normalization. **a** Germination of wild type (Col-8) AR (DAH 554) seeds used for the gene expression analysis. The seeds were imbibed for 7 days. **b** Expression of ABA biosynthesis enzyme genes, *NCED*s, in seeds imbibed for 24 h. **c** Expression of ABA catabolism enzyme genes. The expression of *CYP707A1* and *CYP707A3* were analyzed at 24h after the start of imbibition, and *CYP707A2* was analyzed at 3h after the start of imbibition. **d** Expression of GA biosynthesis enzyme genes, *GA3ox1* and *GA3ox2* in seeds imbibed for 12 h.

GA3ox1 and GA3ox2 are the key enzymes of active GA biosynthesis, and the expression of the genes are regulated by the germination stimulating signals, light and temperature (Toyomasu et al. 1998, Yamaguchi et al. 1998, Yamauchi et al.2004, Toh et al. 2008). Under both in the permissive and non-permissive supra-optimal temperature conditions, *GA3ox1* and *GA3ox2* showed significantly higher expression levels in *MAPKKK20^OX^* seeds than in WT seeds (Fig. 8d). These results suggest the MKK3-MAPK cascade stimulates the expression of GA biosynthesis enzyme genes.

## Discussion

We identified the MAPKKK19/20-MKK3-MPK1/2/7/14 cascade module (MKK3 module) as a mediator of the temperature signal to control Arabidopsis seed germination. The MKK3 module is not required for germination itself since the loss of function alleles of MKK3 could germinate at the optimal temperature condition (Fig. 1c). However, the MKK3 module had a critical role in the modulation of germination temperature range of the seeds, and positively regulated germination at both supra- and sub-optimal temperatures (Fig. 1c, Supplementary Fig. 1f, Fig. 3, Supplementary Fig. 5c, Fig. 5, Supplementary Fig. 8c). The germination temperature range is closely related with dormancy, and expands during after-ripening of winter and summer annual seeds (Baskin & Baskin 2014, Fig. 1c). *MKK3* has been considered as a negative regulator of dormancy in cereals (Torada et al. 2016, Nakamura et al. 2016), but our data suggest that its primary function is as a modulator of germination response to temperature. In addition, the MKK3 module is not essential for after-ripening since the germination temperature range was expanded even in the loss-of-function mutant seeds during dry storage of the seeds (Fig. 1c).

In this study, the MKK3 module activity was regulated by internal and external signals, namely after-ripening of the seeds and environmental temperature (Fig. 9). The MAP kinase cascade can be rapidly activated post-translationally, but relatively slow activation of clade III *MAPKKK*s at the transcriptional level has been reported for ABA, nitrate, and various stress responses (Colcombet et al. 2016). Our data suggest that transcriptional regulation of the two clade III *MAPKKK*s, *MAPKKK19* and *MAPKKK20,* by temperature is key for activation of the MKK3 module and seed germination (Fig. 2, Fig. 6, Fig. 9). This temperature dependent gene expression is modulated by after-ripening, as demonstrated by the fact that repression of *MAPKKK19/20* expression in FH seeds at 26 °C was relieved in AR seeds at the same temperature (Fig. 1a, Fig. 2, Fig. 6, Fig. 9). It has been shown that the germination regulator gene *SOMNUS* is regulated by temperature as well as by light (Lim et al. 2013), but the temperature sensing and signaling mechanism in seeds still needs to be clarified. Thus, elucidation of the temperature signaling and its modulation process during after-ripening may be important for better understanding of seed dormancy and germination. Rice is a summer-annual species, and the germination temperature range expands to downwards during after-ripening, which is in contrast to the winter-annual wheat, barley and Arabidopsis. Recently, it has been suggested that rice MEKK family genes, *OsMAPKKK62* and *OsMAPKKK63* are negative regulators of seed dormancy (Mao et al. 2019, Na et al. 2019). Therefore, it would be interesting to compare the temperature signaling systems between summer and winter annual plants.

**Fig. 9.**
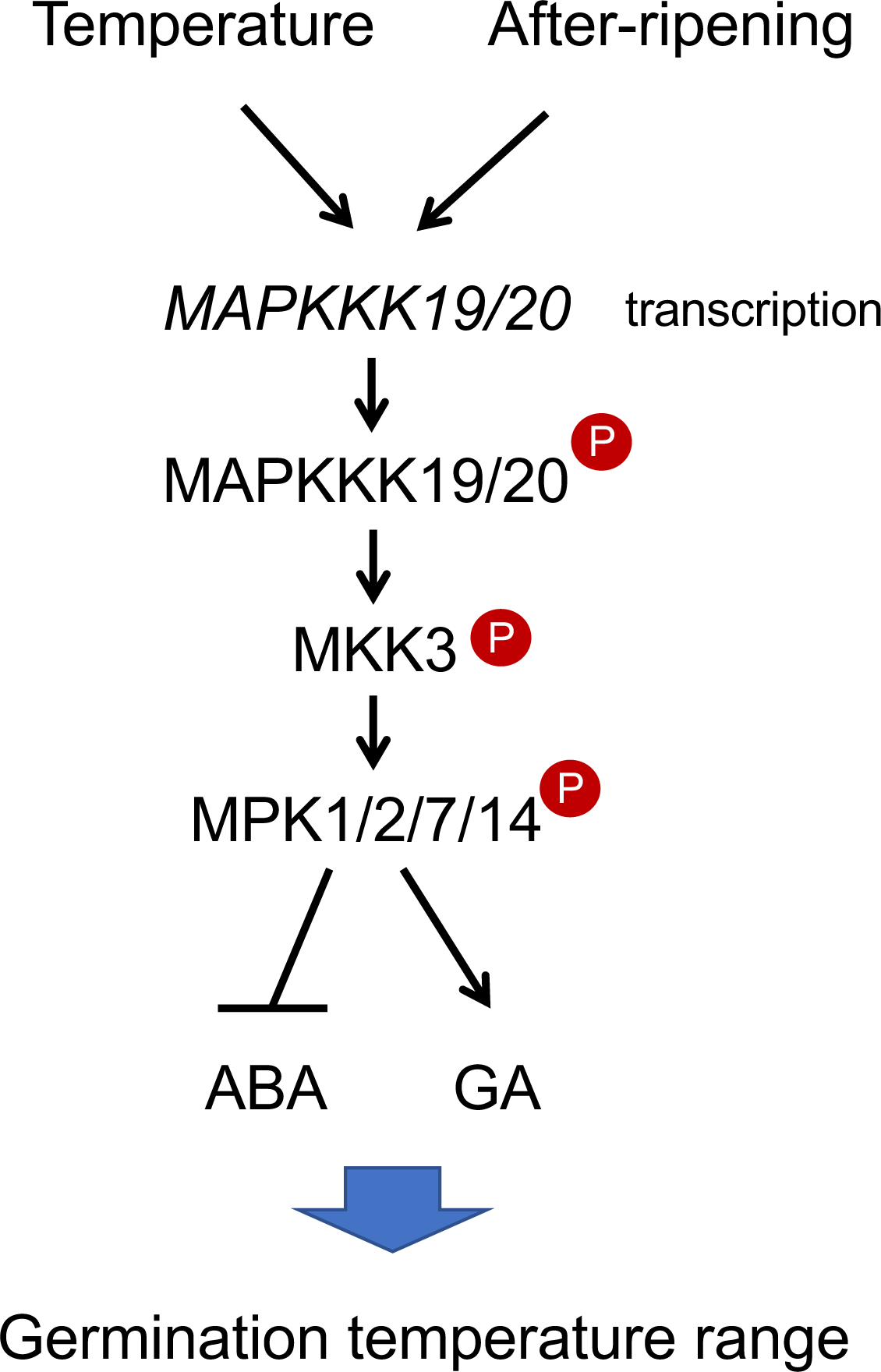
Modulation of germination temperature range by MAPKKK19/20-MKK3-MPK1/2/7/14 module. A model. The MKK3 module is regulated by temperature through the regulation of *MAPKKK19/20* expression, and it regulates seed germination at sub- and supra-optimal temperature conditions through modulation of ABA and GA metabolism. Expression of *MAPKKK19/20* is regulated by temperature and unknown signal of after-ripening. Accumulation of MAPKKK19 and MAPKKK20 proteins induce activation of the MKK3 module, and the kinase cascade signaling decreases ABA accumulation and increases GA production. During after-ripening of the seeds, an unidentified mechanism modulates *MAPKKK19/20* expression, and expands the permissive temperature range of *MAPKKK19/20* induction by suppressing ABA production.

Our data indicate that supra-optimal high temperature represses *MAPKKK19* and *MAPKKK20* expression in the imbibed seeds, and that the MKK3 module regulates seed germination by modulating ABA and GA metabolism (Fig. 7, Fig. 8, Fig. 9). However, in the nitrate induced germination system, the expression of the hormone metabolism genes was not regulated by the MKK3 module (Regnard et al. 2024). In the temperature induced germination system, we could not find any contribution of *MAPKKK13* or *MAPKKK14* (Supplementary Fig. 6), but expression of these genes was induced by nitrate, and they were partially responsible for the activation of MPK7 in response to nitrate (Regnard et al. 2024). These observations suggest that while different environmental signals can activate the MKK3 cascade, there is a signal specific transduction mechanism that controls transcription of specific *MAPKKK*s in the cascade activation.

Our results suggest that the expression of *MAPKKK19/20* is not controlled by ABA, but *MAPKKK17* and *MAPKKK18* have been shown to be induced by ABA in Arabidopsis seedlings, and the MAPKKK17/18-MKK3-MPK1/2/7/14 module has been shown to be involved in the regulation of leaf senescence (Matsuoka et al 2015, Danquah et al 2015). However, in the current study we could not detect any expression of *MAPKKK17/18* in the imbibed seeds, and the seeds of *mapkkk17/18* double mutant showed no germination phenotype (Supplementary Fig. 2, Supplementary Fig. 8). Therefore, tissue, stage and signal specific regulation of different clade III *MAPKKK*s may support the diverse functions of the MKK3 containing MAPK cascade to control plant growth and development.

In addition to MAPKKK19/20, MAPKKK21 was also revealed to be involved in the regulation of FH seed germination. The limit of MAPKKK21, to only having a role in FH seed germination may be explained by its expression only occurring during the seed developmental stage (Fig. 2, Supplemental Fig. 4). MAPKKK21 may activate downstream MKK3 and group C MPKs during seed development, which can modulate the temperature response of freshly matured seeds (Fig. 6c).

In the MKK3 module, among the group C MPKs, MPK7 was shown to have a major role in seed germination, especially at supra-optimal temperatures (Fig. 5, Supplementary Fig. 8). We also found that MPK2 had a major contribution to germination at sub-optimal low temperatures (Fig. 5c, Supplemental Fig. 11c). Therefore, different group C MPKs may have different substrate specificity, and different systems may regulate seed germination at supra- and sub-optimal temperature conditions.

MPK7 activity showed dual peaks at 3 h and from 12 to 24 h after the start of imbibition during germination (Fig. 6). The first activity peak may not be sufficient for the completion of germination since this initial activity was also detected in non-germinating after-ripened seeds imbibed at supra-optimal temperature. The second MPK7 activity peak was not observed at non-permissive supra-optimal temperatures in either FH or AR seeds, suggesting that it has a critical role for germination (Fig. 2, Fig. 6, Fig. 9). During the germination process, seed water uptake is divided in to three phases, passive and rapid water uptake (phase I), stationary (phase II) and seedling growth associated water uptake (phase III) (Bewley 1997). In Arabidopsis, phase I is completed by 1 to 3 h after the start of imbibition, and radicle protrusion is observed after around 30 h (Preston et al. 2009). Therefore, the MPK7 activity peaks observed in the current study correspond to the early and late stages of phase II. In the early stage of phase II, the MKK3 module may target pre-existing and newly synthesized proteins which stimulate the initiation of the germination process in AR seeds, including respiration, macromolecule repair, transcription and translation (Bewley et al. 2013, Preston et al. 2009). In the late stage of phase II, the MKK3 module may phosphorylate proteins which stimulate the completion of the germination process including ABA and GA metabolism.

Collectively, the MKK3 module integrates internal after-ripening and external temperature signals into the germination regulation process, and modulates the germination temperature range which is critical to establish the dormancy levels and germination timing of the seeds.

## Materials and Methods

### Plant materials and growth conditions

T-DNA insertion lines of Arabidopsis (*Arabidopsis thaliana* L. Heynh.) Columbia-0 (Col-0) accession were obtained from the Arabidopsis Biological Resource Center [*mkk3-1* (SALK_051970; Takahashi *et al*., 2007), *mkk3-2* (SALK_208528C; Sözen *et al*., 2020), *mpk1-1* (SALK_063847C; Enders *et al*., 2017), *mpk2-2* (SALK_047422C; Lv *et al*., 2021), *mpk7-1* (SALK_113631), *mpk14-1* (SALK_022928C; Lv *et al*., 2021), *mapkkk13-1* (GK-277E09), *map3k14-1* (GK-653B01; Sözen *et al*., 2020), *mapkkk15-1* (SALK_084817), *map3k16* (Choi *et al*., 2017), *mapkkk17-2* (SALK_080309C; Romero-Hernandez and Martinez, 2022), *mkkk18-2* (GK-676E02; Mitula *et al*., 2015) and *mapkkk20-3* (GK-458D07)]. Ds-transposon insertion lines of Arabidopsis Nossen accession were obtained from RIKEN BioResource Research Center [*mapkkk19-1* (pst14411) and *mapkkk21-3* (psh20310)]. Homozygous insertion lines were selected by PCR with the specific primer sets (Supplementary Table 1 and 2). Genome editing lines of Arabidopsis Col-0 accession *mapkkk13/14CR* were created by using the CRISPR-Cas9 system (Sözen et al. 2020, Regnard et al. 2024). *MAPKKK20^ox^* lines were created by transformation of Col-8 accession using the pGWB20 destination vector (Nakagawa et al. 2007). *aba2-2* (Nambara *et al*., 1998) was kindly provided by Dr. E. Nambara (Toronto University, Toronto), and *ga3ox1-3* (SALK_004521) and *ga3ox1-3 ga3ox2-1* were kindly provided by Dr. E. Nambara (Toronto University, Toronto) and Dr. S. Yamaguchi (Kyoto University, Kyoto), respectively (Nambara et al. 1998, Mitchum et al. 2006).

To generate multiple group C *MPK* mutants, we first isolated *mpk1-1 mpk2-2* and *mpk7-1 mpk14-1* double mutants. Then, we crossed the double mutants, and isolated quadruple, triple, double, single mutants and the wild type siblings from the segregants by PCR, as described above.

*mapkkk19-1* and *mapkkkk21-3* in Nossen background were backcrossed four times with Col-0, and the introgression lines *mapkkk19-1C* and *mapkkk21-3C* were selected from BC_4_F_2_ and BC_3_F_2_ siblings, respectively, by PCR. To obtain multiple *mapkkk* mutants, we first crossed *mapkkk19-1C* and *mapkkk21-3C*, and then crossed the F_1_ plant with *mapkkk20-3*. Then, multiple mutants were isolated from F_2_ and F_3_ plants. Genotyping was done by PCR with the gene-specific primers and either T-DNA left-border or Ds-transposon H-edge primers (Supplementary Table 2). We crossed *MAPKKK20^OX^* (#125) and *mkk3-1* and selected double mutants from the F_2_ plants by PCR using specific primer sets for the overexpression vector pGWB20 and T-DNA insertion in *MKK3* (Supplementary Table 2).

Seeds were stratified for 4 days at 4 °C and then directly sown and grown in soil (Co-op N-150; Katakura & Co-op Agri, Co., Tokyo, Japan) in a growth chamber (continuous illumination at 22 °C). Then, after the plants reached physiological maturity, when about one-half of the fruits on a plant turned to yellow, their seeds were harvested and dried for 2 days in a desiccator. These were used as freshly harvested (FH) seeds. After-ripened (AR) seeds were obtained by storing FH seeds in a desiccator at room temperature for at least 2 months. To maintain dormancy, some of the FH seeds were stored at −80 °C with silica gel.

### Germination assay

Thirty seeds were imbibed with 300 μL of ultra-pure water in the well of a 24-well plate at constant temperature in continuous light for 5 to 14 days without cold stratification, unless otherwise stated. Germination was scored as radicle protrusion. The germination ability of the seeds was evaluated as the final germination percentage or as the Germination Index (GI). The GI was calculated with maximum weight given to the seeds that germinated early and less weight given to those that germinated late as follows: 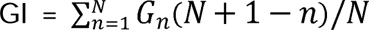, where *G_n_* is the percentage of germinated seeds on day *n*, but not a cumulative value. Germination tests were done with at least three independent seed batches with three replicates in each, unless otherwise stated. Each batch contained the seeds harvested from at least 4 plants. A typical result was presented because there was some variation in germination percentages between batches, but the relative differences between treatments in any batch were very similar for all batches.

For the chemical treatments, 30 seeds were imbibed with 250 μL of hormone solutions in the well of a 24-well plate at constant temperature in continuous light for 5 to 7 days without cold stratification. ±ABA (SIGMA, A1049), GA_3_ (SIGMA, G7645) and fluridone (Daw Elanco, recrystallized) were first dissolved in dimethylsulfoxide (DMSO) and then diluted to the final concentrations with ultra-pure water. The final concentration of DMSO was 0.1%, and 0.1% DMSO solution was used as the control.

### RNA isolation and qRT-PCR analysis

Dry or imbibed seeds (15-20 mg dry weight), seedlings and siliques were frozen with a φ 5mm stainless bead in LN_2_ immediately after sampling, and stored at -80 °C until use. The seeds in developmental stages were collected as described previously (Zheng et al. 2022). In brief, we marked open flowers at anthesis (day zero) by thread, and collected the fruits at 3, 6, 9, 12, 15 and 18 days post-anthesis (DPA). Total RNAs were isolated from 20 siliques with seeds (DPA 3 and 6) and seeds collected by opening the 20 siliques (DPA 9 to 18). The frozen tissues were squashed in a 2 ml tube by a bead mill type homogenizer (Biomedical Science, Tokyo). Total RNA was extracted using the hexadecyltrimetylammonium bromide method as described previously (Zheng et al. 2022). RNA was reverse-transcribed to cDNA by using DNA removal and a cDNA synthesis kit (PrimeScriptTM RT reagent Kit with gDNA Eraser; TaKaRa, Kusatsu, Japan) with a mixture of oligo-dT and random primers. Quantification of transcript was done by qRT-PCR with fluorescent-labelled nucleotide substrate (TB Green™ Premix Ex Taq™ II, Takara or PowerUp SYBR Green Master Mix, Thermo Scientific) as described previously (Shigeyama et al. 2016, Zheng et al. 2022). Forward and reverse primer sequences for semi-quantitative RT-PCR and qRT-PCR are listed in Supplementary Table 2. Reactions were done using the 7500 Fast system (Applied Biosystems, ABI), and the data were analyzed using ABI Prism 7700 SDS software (ABI). For each sample, the mean value from triplicate qRT-PCRs was used to calculate the transcript abundance. At2g20000 was used as a reference genes for transcript normalization (Graeber et al. 2011). For each sample, the mean value from triplicate reactions was used to calculate transcript abundance, with the mean values being plotted along with the standard deviations. To confirm biological reproducibility, experiments were performed at least three times using samples harvested in different batches; similar results were obtained. Typical results are presented unless otherwise stated.

### Microarray analysis

Freshly harvested Col-0 seeds were imbibed at 26 oC, and after-ripened seeds were imbibed at either 26 °C or 34 °C under continuous illumination. Total RNA was extracted from 20 mg seeds (dry weight) using the RNAqueous™ Small scale phenol-free total RNA isolation kit (Ambion: catalog #1912) according to the manufacturer’s protocol. Three seed batches harvested from independently grown plants were used for biological replications. Cyanine 3-labelled cRNA was synthesized from 150 ng RNA using the Low Input Quick Amp Labeling Kit (Agilent) and predicated using RNeasy Mini Kit (QIAGEN) according to the manufacturer’s protocol. The labeled cRNA was fragmented and hybridized to Agilent Arabidopsis 4 Oligo Microarrays (G2519F) for 17 h at 65 oC. After hybridization on 4 x 44K array slide, the arrays were washed and scanned by Agilent DNA Microarray Scanner (G2505B) according to one-color methods. Signal intensities were measured by Feature Extraction Software 11.5.1.1 (Agilent) and data analysis were performed by Gene Spring (Agilent) and R software. Normalization was conducted using Modified Histogram Matching Normalization (MHMN) method (Astola and Molenaar, 2014) . Araport11 (TAIR) was set for gene annotation. Raw data is available at GEO database (Clough and Barrett 2016); accession #GSE229182.

### Kinase assay of MPK7 in seeds

FH and AR seeds (10 mg) were imbibed at permissive or supra-optimal temperatures for germination under continuous illumination. The seeds were frozen in LN_2_ immediately after sampling, and stored at -80 °C until use. Protein extraction, kinase assay and western blotting were performed as described (Sözen et al. 2020, Regnard et al. 2024). In brief, frozen seeds were ground, and then soluble proteins were extracted using a non-denaturing buffer, supplemented with phosphatase inhibitors. After normalization on total protein amount, MPK7 was immunoprecipitated using a specific antibody, and its activity was assayed as the ability to phosphorylate the substrate MBP. Phosphorylated MBP was revealed on a SDS-PAGE gel. Western blots were performed using indicated antibodies (Sözen et al. 2020, Regnard et al. 2024).

## Acknowledgments

We thank Dr. Atsushi Torada at Hokuren and Dr. Taishi Umezawa at Tokyo University of Agriculture and Technology greatly for valuable discussion. We thank Dr. Tsuyoshi Nakagawa in Interdisciplinary Center for Science Research, Shimane University for providing pGWB vector plasmids. We also thank Dr. Eiji Nambara at Toronto University and Dr. Shinjiro Yamaguchi at Kyoto University for providing ABA deficient (*aba2-2*) and GA deficient (*ga3ox1* and *ga3ox2*) mutant seeds, respectively. We thank RIKEN BioResource Research Center and ABRC for providing transposon and T-DNA insertion mutant seeds, respectively. We thank Kohei Yokota at Faculty of Agriculture, Kagawa University, Yuichi Kashiwakura and Ryoshun Nishioka at the Department of Life Sciences, Meiji University, for their technical assistance. We thank Dr. Iain McTaggart at School of Agriculture, Meiji University for critical reading and English editing of the manuscript. This work was supported partly by the Program for the Strategic Research Foundation at Private Universities, 2014–2018 of MEXT, Japan.

## Author contributions

M.O. and N.K. conceived the project; M.O. and N.K. conducted experiments and data analysis; M.O., R.T., L.Z., T.H. and S.O. performed mutant isolation and characterization; S.R. performed kinase assays; K.I. constructed *MAPKKK20^ox^*lines; T.S. preformed microarray experiment; N.T. and S.K. performed the bioinformatics; J.C. and K.I. contributed to discussion; M.O. wrote the initial manuscript; All authors edited the manuscript.

## Data availability

The microarray data generated from this study have been deposited in the Gene Expression Omnibus under accession code GSE229182. The unique biological materials are available upon appropriate requests. Source data are provided with this paper.

## Legends to Supplementary Figures

**Supplementary Fig. 1 Germination of *MKK3* loss-of-function mutant seeds.**

(**a**) Gene model of *MKK3* and the position of T-DNA insertion. (**b, c**) Germination time course of WT (Col-0), *mkk3-1* and *mkk3-2* FH (DAH 2) seeds. The seeds were imbibed at 22 °C without stratification. Typical germination time course data from three (b) or two (a) biological replicates are shown. The values are the means (±SD) of three technical replicates, and we had similar results in different replicates. (**d, e**) Germination time course of FH and AR seeds imbibed at 5 °C (**d**) and at 10 °C (**e**). The values are the means (±SD) of three biological replicates.

**Supplementary Fig. 2 Expression of clade-III *MEKK*s during imbibition of FH and AR seeds.**

(**a**) Phylogenetic tree of MEKK-like MAPKKK in Arabidopsis. Phylogenetic analysis with amino acid sequences was done by neighbor-joining method using MEGA X (Kumar et al. 2018). Percentages of clustering reproducibility (bootstrap test with 1000 replicates) are shown at the branch points. The evolutionary distances were calculated using the Poisson correction method. (**b**) Expression of clade-III MEKK-like *MAPKKK* during imbibition of dry mature seeds (GEO accession: GSE229182). Values shown are means (±SD) of three biological replicates of microarray analysis with Arabidopsis 4 Oligo Microarray (Agilent).

**Supplementary Fig. 3 T-DNA insertion alleles of *MAPKKK19, 20* and *21*.**

(**a**) Position of T-DNA (white triangles) and transposon (black triangles) insertion in Col-0 (*mapkkk20-3*) and Nossen (*mapkkk19-1* and *mapkkk21-3*) accessions, respectively. Positions of primers used for genotyping (white arrows) and expression (black arrows) analyses are indicated. (**b**) Semi-quantitative RT-PCR for the mutant allele expression analysis. Total RNA was extracted from 24 h imbibed seeds for *MAPKKK19* and *MAPKKK20*, and from dry seeds for *MAPKKK21*. 18s rRNA was used as an internal control. PCR cycle numbers were 27 for *MAPKKK19* and *MAPKKK20*, 30 for *MAPKKK21,* 21 for 18S rRNA.

**Supplementary Fig. 4 Expression of *MAPKKK19/20/21* during seed development.**

Total RNA was prepared from seeds with siliques at 3 and 6 days after flowering (DAF) and from seeds without siliques at 9 to 21 DAF. Transcript levels were quantified by quantitative RT-PCR using At2g20000 as an internal control. The quantification was done with three independent plant populations, and typical data are presented. We obtained similar results from the different experiments.

**Supplementary Fig. 5 Seed dormancy and germination of *mapkkk19/20/21* at sub-optimal temperature.**

Asterisks indicate statistical differences between wild type (Col-0) and *mapkkk19/20/21* (p < 0.05, Student’s t test). (**a**) Enhanced dormancy phenotype of *mapkkk19/20/21* seeds. Freshly harvested seeds were stored in a desiccator for up to 3 weeks at room temperature. Seeds were imbibed at 22 °C for 7 days without cold stratification. Values are means (±SD) of three technical replicates. We obtained similar results from triplicate experiments, and typical data are presented. (**b**) Effect of cold stratification on germination of the freshly harvested seeds. The seeds at DAH 2 were imbibed at 22 °C for 5 days with (+) or without (−) preceding cold stratification at 4 °C for 4 days. (**c**) Germination time course of AR seeds imbibed at 5 °C. (**b, c**) Results from three independent seed batches are shown with averages and SDs.

**Supplementary Fig. 6 Germination of gain- and loss-of-function mutant seeds of *MAPKKK13* and *MAPKKK14*.**

(**a**) Gene model of *MAPKKK13* and *MAPKKK14*, and the positions of T-DNA insertion (*mapkkk13-1, mapkkk14-1*) and gene editing (*mapkkk13CR, mapkkk14CR*) positions. Asterisk indicates stop codon created by single nucleotide insertion by CRISPR-Cas9. (**b**) Semi-quantitative RT-PCR for *mapkkk13-1* expression analysis. Total RNA was extracted from dry seeds. 18s rRNA was used as an internal control. PCR cycle numbers were described above the gel image. (**c**) Germination of the gain-of-function mutant seeds. FH (DAH 2) and AR (DAH 111-113) seeds were imbibed for 7 days. We obtained similar results from triplicate experiments, and typical data are presented. The values were presented as mean (±SD) of three technical replicate. (**d**) Germination of loss-of-function *mapkkk13/14-cr1* and *mapkkk13/14-cr2* double mutant seeds. FH (DAH 2) were imbibed for 7 days, and AR (DAH 390) seeds were imbibed for 5 days. Results from three independent seed batches are shown with averages and SDs.

**Supplementary Fig. 7 Germination of loss-of-function mutant seeds of *MAPKKK15* and *MAPKKK16*.**

(**a**) Gene model of *MAPKKK15* and *MAPKKK16*, and the positions of T-DNA insertion positions. (**b**) Semi-quantitative RT-PCR for the mutant gene expression analysis. Total RNA was extracted from 7-days-old seedlings treated with 10 µM ABA for 1 h. 18s rRNA was used as an internal control. PCR cycle numbers were 30 for *MAPKKK15* and *MAPKKK16*, and 21 for 18S rRNA. (**c**) Germination of FH (DAH 2) and AR seeds (DAH 111-113). The seeds were imbibed for 7 days. The experiments were performed in three independent seed batches with three replicates in each. We obtained similar results from the three experiments, and typical data are presented. The values were presented as mean (±SD) of three technical replicates (n = 8). We could not find significant differences between WT and the mutants (Tukey HSD tests, PL<L0.05).

**Supplementary Fig. 8 Germination of loss-of-function mutant seeds of *MAPKKK17* and *MAPKKK18*.**

(**a**) Gene model of *MAPKKK17* and *MAPKKK18,* and the positions of T-DNA insertion positions. Positions of primers used for expression analysis are indicated by arrows. (**b**) Semi-quantitative RT-PCR for the mutant gene expression analysis. Total RNA was extracted from 7-day-old seedlings treated with 10 µM ABA for 3 h. 18s rRNA was used as an internal control. PCR cycle numbers were 30 for *MAPKKK17* and *MAPKKK18*, and 21 for 18S rRNA. (**c**) FH (DAH 2) and AR (DAH 80) seeds were imbibed for 7 and 5 days, respectively. We obtained similar results from triplicate experiments, and typical data are presented. We could not find significant differences between WT and the mutants (Tukey HSD tests, PL<L0.05).

**Supplementary Fig. 9 Over-expression of *MAPKKK20* and the effect on germination at sub-optimal temperature.**

(**a**) Schematic representation of *MAPKKK20^OX^* construct. Positions of primers used for expression analysis are indicated by arrows. (**b**) Quantification of *MAPKKK20* transcripts by qRT-PCR with At2g20000 as an internal control. Relative expression to WT (Col-8) is indicated by the fold change expression in *MAPKKK20^OX^* dry seeds. The values are the means (±SD) of three technical replicates. (**c**) Germination time course of AR (DAH 167) seeds imbibed at 5 oC. The values are the mean (±SD) of three biological replicates.

**Supplementary Fig. 10 T-DNA insertion alleles of group C MPKs.**

(**a**) Schematic representation of the genes and position of T-DNA (white triangles) insertion in Col-0. Positions of primers used for genotyping (white arrows) and expression (black arrows) analyses are indicated. (**b**-**d**) Semi-quantitative RT-PCR for the mutant allele expression analysis. Total RNA was extracted from 24 h imbibed seeds. 18s rRNA was used as an internal control. PCR cycle numbers were 30 for *MPK*s and 21 for 18S rRNA.

**Supplementary Fig. 11 Seed dormancy and germination of group C MPK multiple mutants at sub-optimal temperature.**

(**a**) Enhanced dormancy phenotype of *mpk1/7/14 and mpk1/2/7/14* seeds. Freshly harvested seeds were stored in a desiccator at room temperature. Seeds were imbibed at 22 °C for 7 days without cold stratification. Values are means of three technical replicates with SDs. We obtained similar results from triplicate experiments, and typical data are presented. (**b**) Effect of cold stratification on germination of the freshly harvested seeds. The seeds at DAH 2 were imbibed at 22 °C for 5 days with (+) or without (−) preceding cold stratification at 4 °C for 4 days. (**c**) Germination time course of AR seeds imbibed at 5 °C. (**a, b**) Significant differences between samples are indicated by different letters (Tukey HSD tests, PL<L0.05). (**b**, **c**) Results from three independent seed batches are shown with averages and SDs.

**Supplementary Fig. 12 MPK7 activity in germinating and non-germinating seeds (Biological replication of Fig. 6).** In panel c, electrophoresis and staining of the proteins in the kinase assay mixture has not been done.

## Supplementary Data

**Supplementary Table 1** Genes and its mutant alleles used in this study

**Supplementary Table 2** Primers used in this study

