## Supplemental Figures and Tables for "The MKK3 MAPK cascade integrates temperature and after-ripening signals to modulate seed germination"

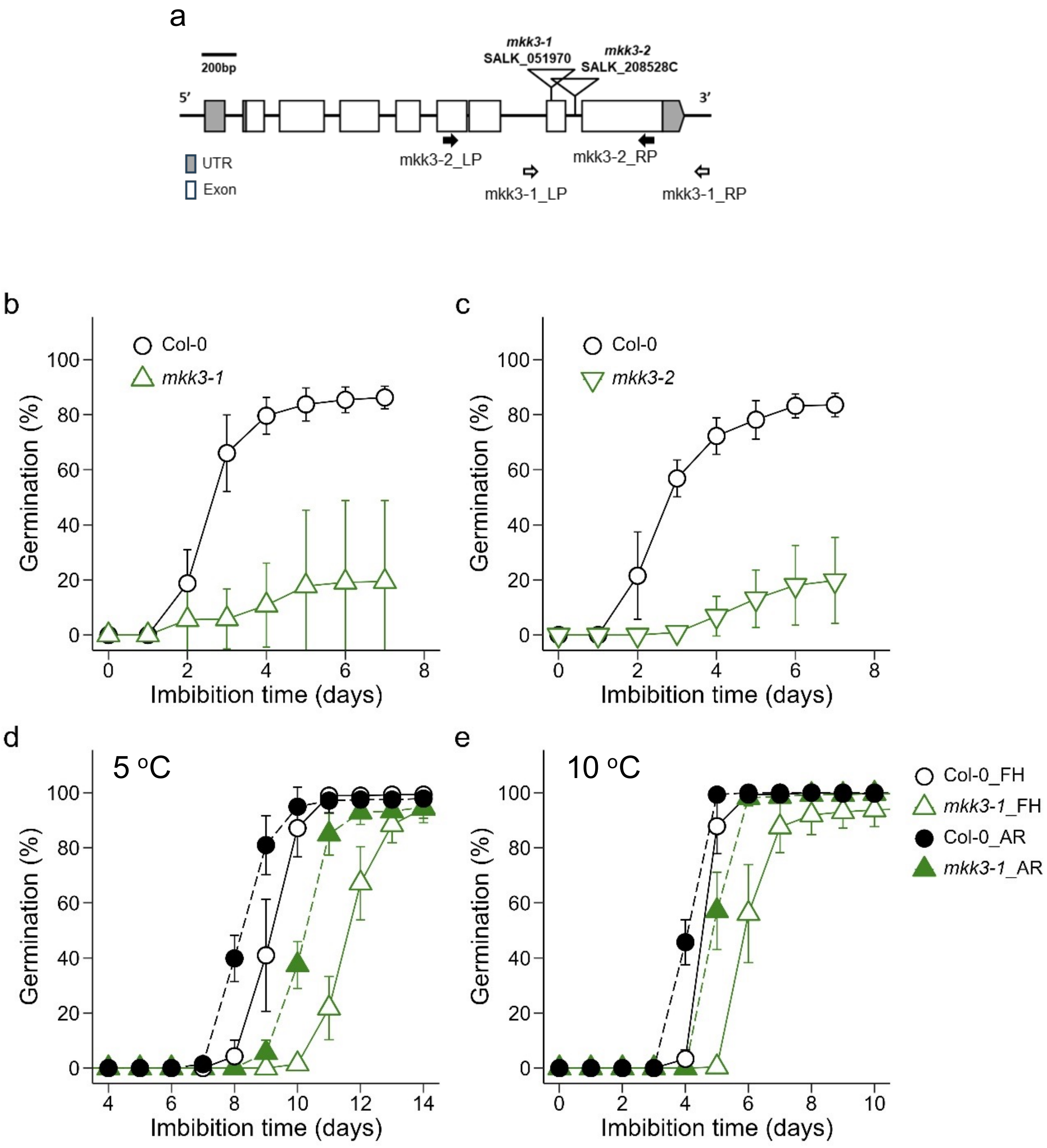


**Supplementary Fig. 1 Germination of *MKK3* loss-of-function mutant seeds.**

(**a**) Gene model of *MKK3* and the position of T-DNA insertion. (**b, c**) Germination time course of WT (Col-0), *mkk3-1* and *mkk3-2* FH (DAH 2) seeds. The seeds were imbibed at 22 °C without stratification. Typical germination time course data from three (b) or two (a) biological replicates are shown. The values are the means (±SD) of three technical replicates, and we had similar results in different replicates. (**d, e**) Germination time course of FH and AR seeds imbibed at 5 °C (**d**) and at 10 °C (**e**). The values are the means (±SD) of three biological replicates.

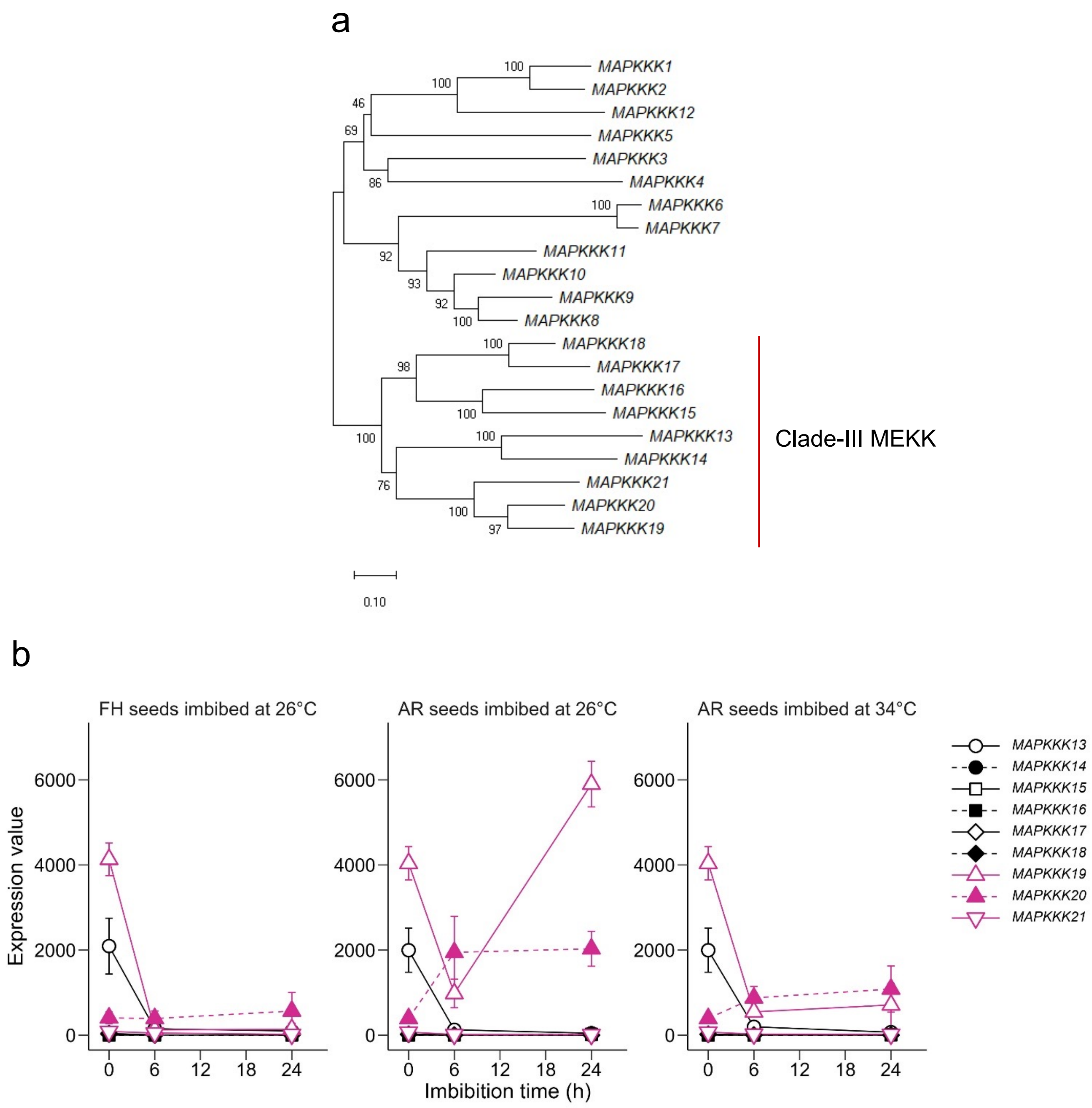


**Supplementary Fig. 2 Expression of clade-III *MEKK*s during imbibition of FH and AR seeds**

(**a**) Phylogenetic tree of MEKK-like MAPKKK in Arabidopsis. Phylogenetic analysis with amino acid sequences was done by neighbor-joining method using MEGA X (Kumar et al. 2018). Percentages of clustering reproducibility (bootstrap test with 1000 replicates) are shown at the branch points. The evolutionary distances were calculated using the Poisson correction method. (**b**) Expression of clade-III MEKK-like *MAPKKK* during imbibition of dry mature seeds (GEO accession: GSE229182). Values shown are means (±SD) of three biological replicates of microarray analysis with Arabidopsis 4 Oligo Microarray (Agilent).

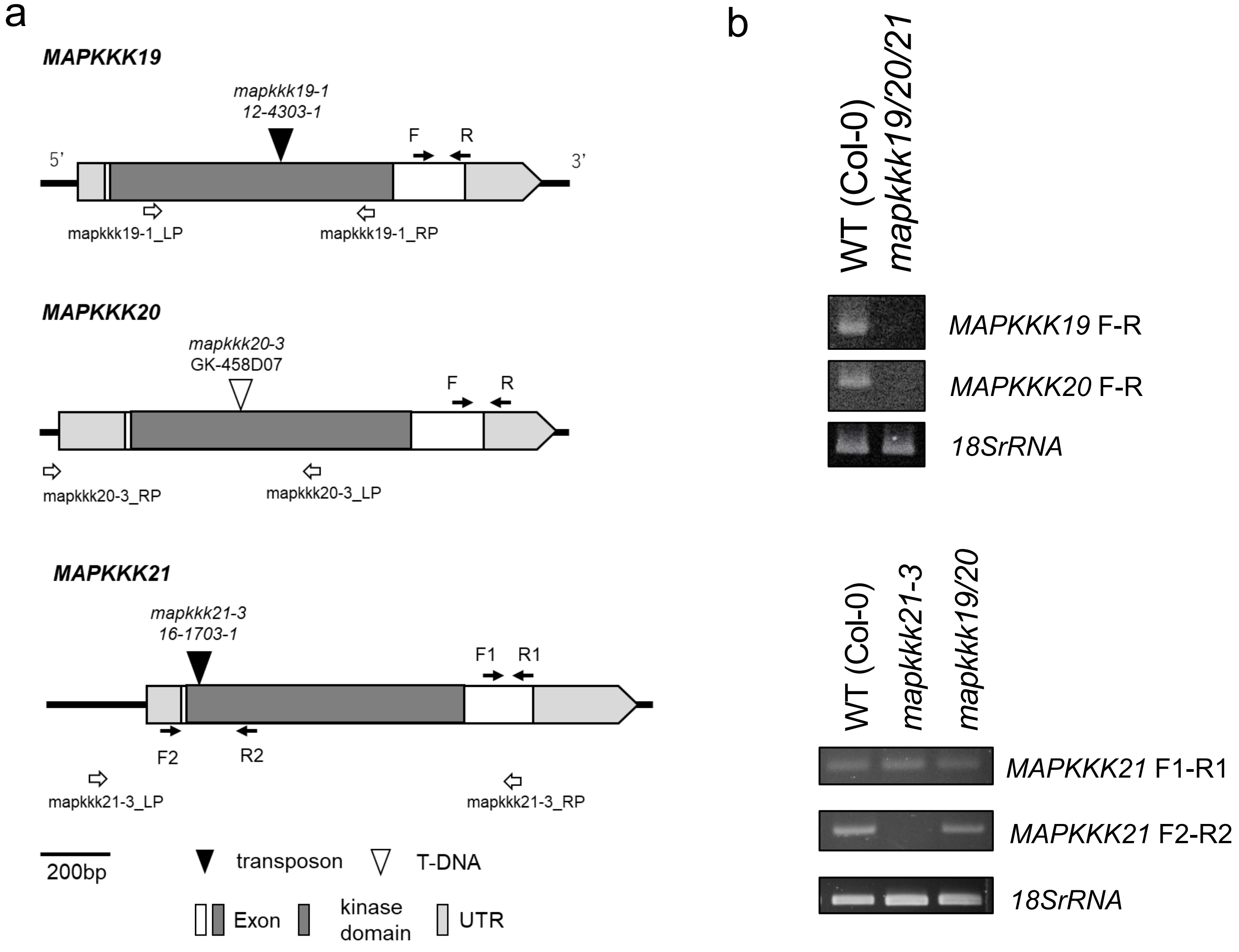


**Supplementary Fig. 3 T-DNA insertion alleles of *MAPKKK19, 20* and *21*.** (**a**) Position of T-DNA (white triangles) and transposon (black triangles) insertion in Col-0 (*mapkkk20-3*) and Nossen (*mapkkk19-1* and *mapkkk21-3*) accessions, respectively. Positions of primers used for genotyping (white arrows) and expression (black arrows) analyses are indicated. (**b**) Semi-quantitative RT-PCR for the mutant allele expression analysis. Total RNA was extracted from 24 h imbibed seeds for *MAPKKK19* and *MAPKKK20*, and from dry seeds for *MAPKKK21*. 18s rRNA was used as an internal control. PCR cycle numbers were 27 for *MAPKKK19* and *MAPKKK20*, 30 for *MAPKKK21,* 21 for 18S rRNA.

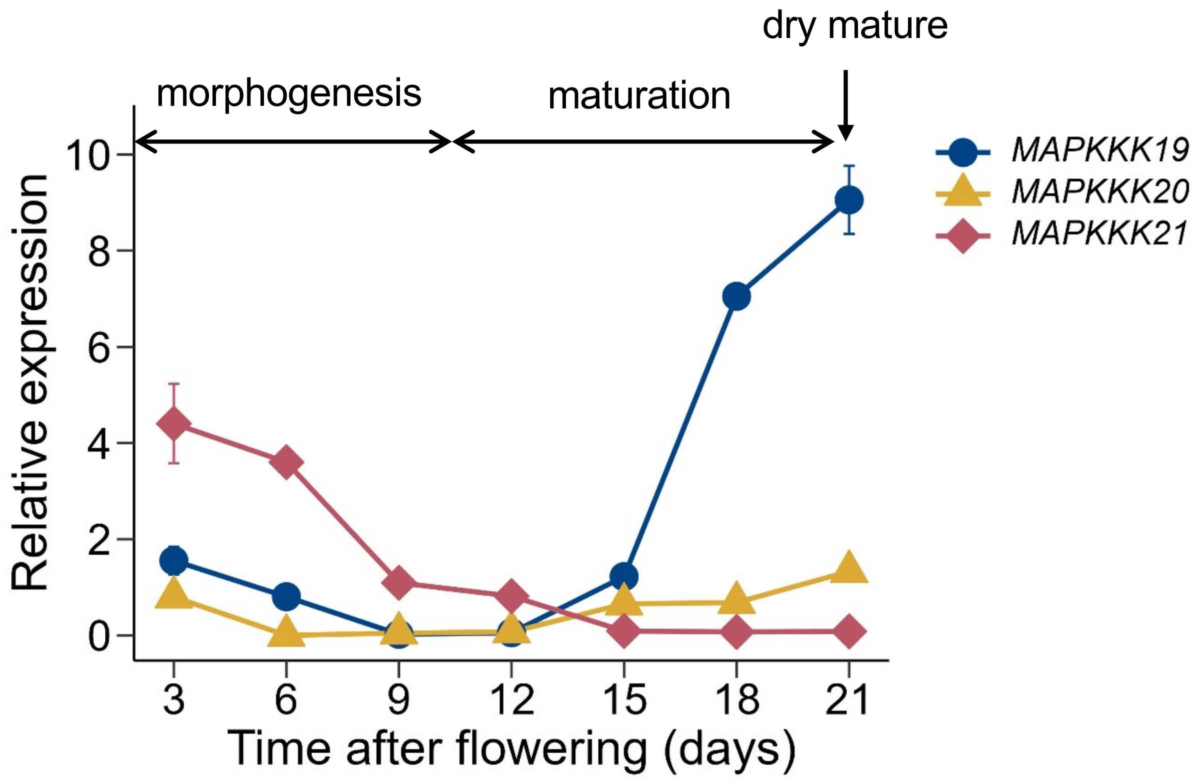


**Supplementary Fig. 4 Expression of *MAPKKK19/20/21* during seed development.**
Total RNA was prepared from seeds with siliques at 3 and 6 days after flowering (DAF) and from seeds without siliques at 9 to 21 DAF. Transcript levels were quantified by quantitative RT-PCR using At2g20000 as an internal control. The quantification was done with three independent plant populations, and typical data are presented. We obtained similar results from the different experiments.

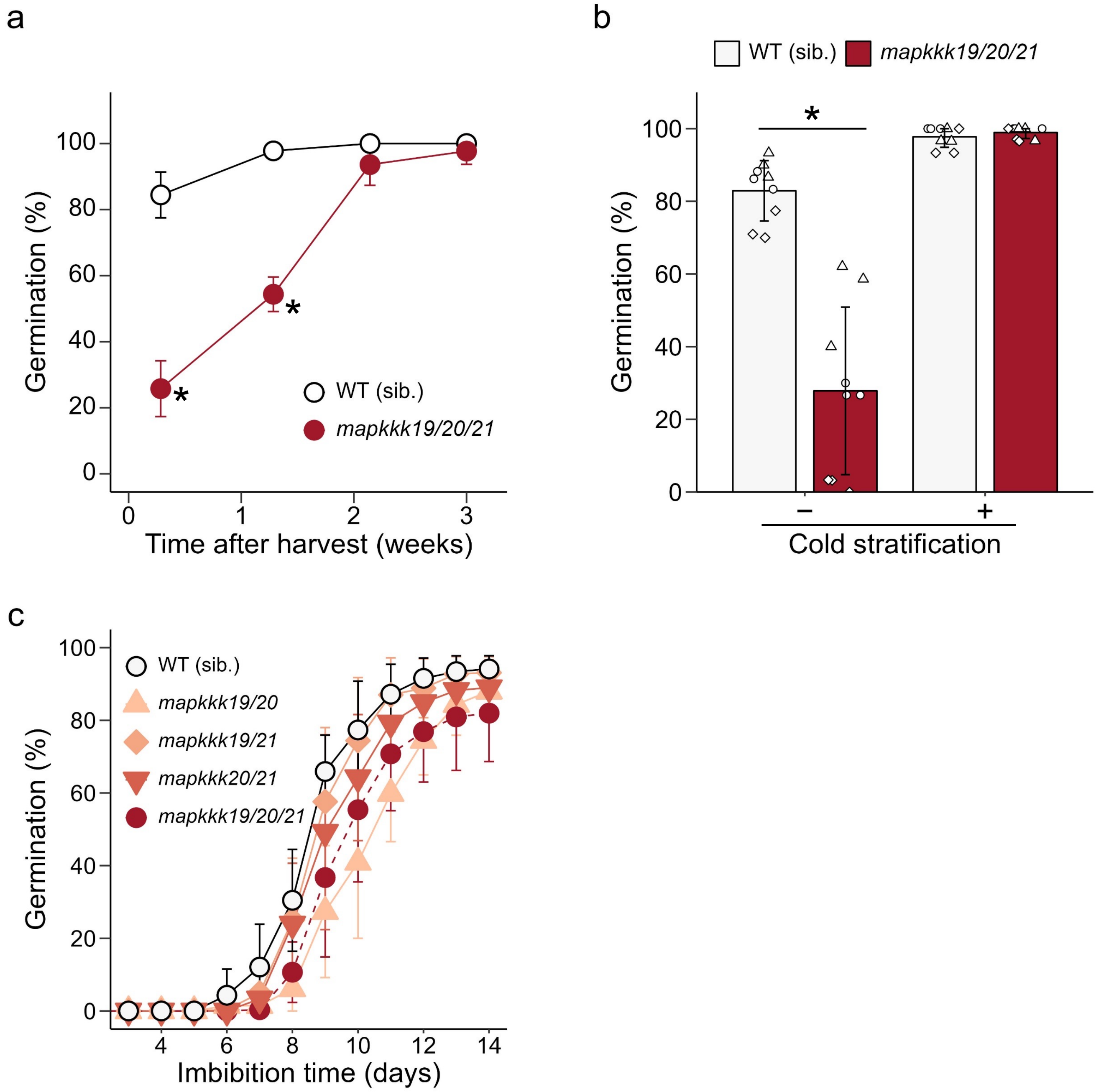


**Supplementary Fig. 5 Seed dormancy and germination of *mapkkk19/20/21* at sub-optimal temperature**

Asterisks indicate statistical differences between wild type (Col-0) and *mapkkk19/20/21* ($p < 0.05$, Student's t test). (**a**) Enhanced dormancy phenotype of *mapkkk19/20/21* seeds. Freshly harvested seeds were stored in a desiccator for up to 3 weeks at room temperature. Seeds were imbibed at 22 °C for 7 days without cold stratification. Values are means (±SD) of three technical replicates. We obtained similar results from triplicate experiments, and typical data are presented. (**b**) Effect of cold stratification on germination of the freshly harvested seeds. The seeds at DAH 2 were imbibed at 22 °C for 5 days with (+) or without (−) preceding cold stratification at 4 °C for 4 days. (**c**) Germination time course of AR seeds imbibed at 5 °C. (**b, c**) Results from three independent seed batches are shown with averages and SDs.

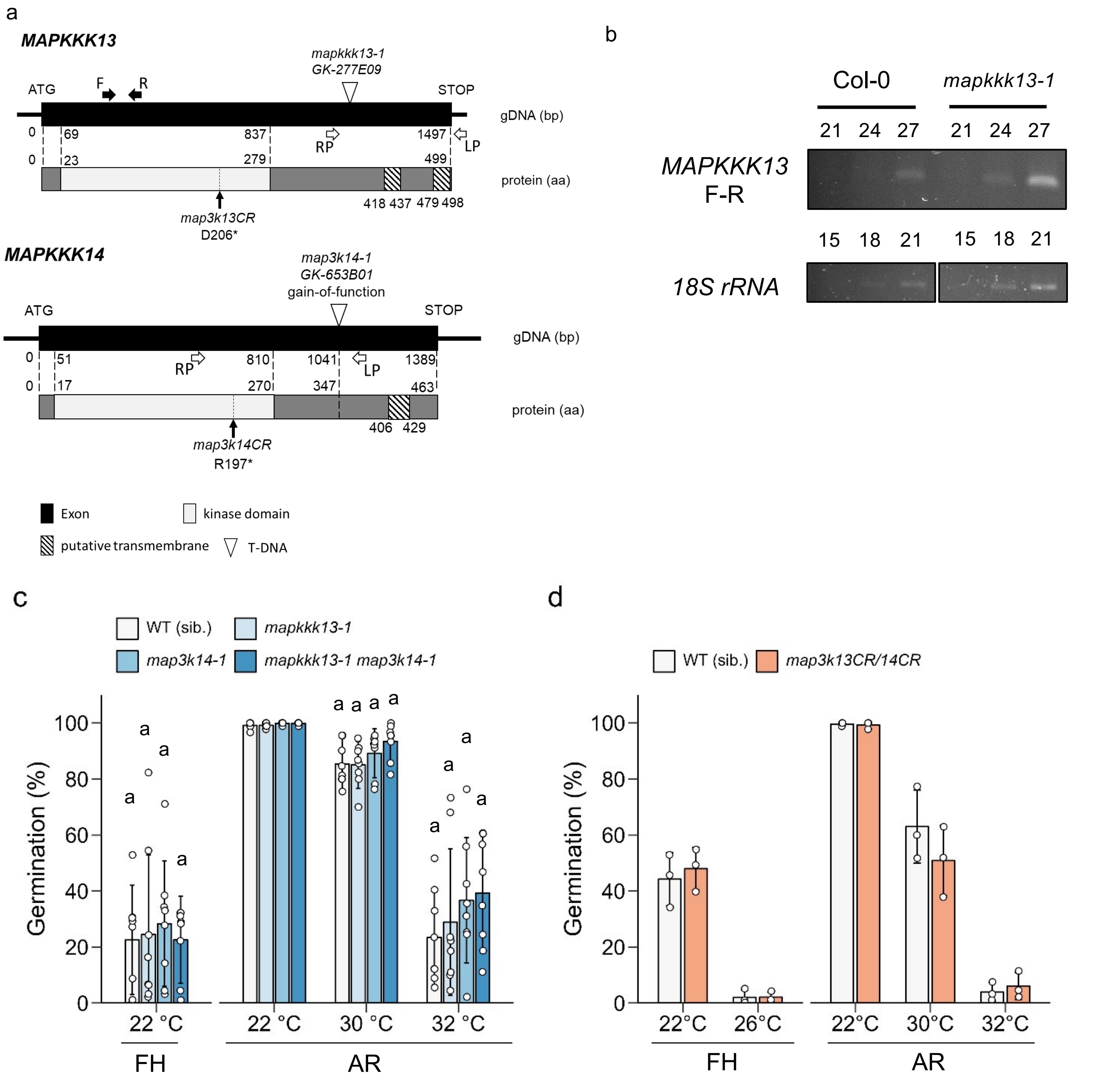


**Supplementary Fig. 6 Germination of gain- and loss-of-function mutant seeds of *MAPKKK13* and *MAPKKK14*.**

(**a**) Gene model of *MAPKKK13* and *MAPKKK14*, and the positions of T-DNA insertion (*mapkkk13-1, mapkkk14-1*) and gene editing (*mapkkk13CR, mapkkk14CR*) positions. Asterisk indicates stop codon created by single nucleotide insertion by CRISPR-Cas9. (**b**) Semi-quantitative RT-PCR for *mapkkk13-1* expression analysis. Total RNA was extracted from dry seeds. 18s rRNA was used as an internal control. PCR cycle numbers were described above the gel image. (**c**) Germination of the gain-of-function mutant seeds. FH (DAH 2) and AR (DAH 111-113) seeds were imbibed for 7 days. We obtained similar results from triplicate experiments, and typical data are presented. The values were presented as mean (±SD) of three technical replicate. (**d**) Germination of loss-of-function *mapkkk13/14-cr1* and *mapkkk13/14-cr2* double mutant seeds. FH (DAH 2) were imbibed for 7 days, and AR (DAH 390) seeds were imbibed for 5 days. Results from three independent seed batches are shown with averages and SDs.

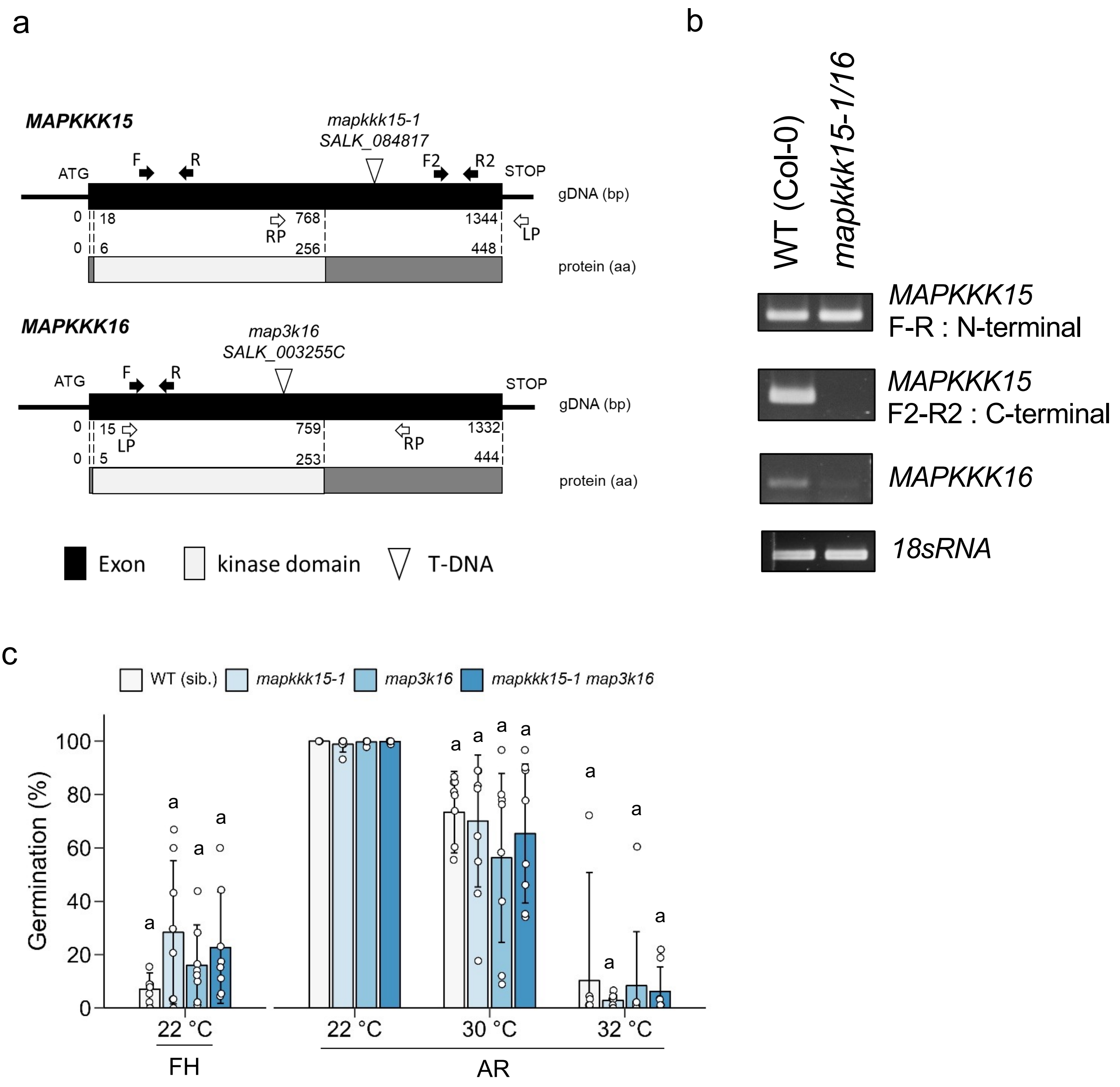


**Supplementary Fig. 7 Germination of loss-of-function mutant seeds of *MAPKKK15* and *MAPKKK16*.** (**a**) Gene model of *MAPKKK15* and *MAPKKK16*, and the positions of T-DNA insertion positions. (**b**) Semi-quantitative RT-PCR for the mutant gene expression analysis. Total RNA was extracted from 7-days-old seedlings treated with 10 μM ABA for 1 h. 18s rRNA was used as an internal control. PCR cycle numbers were 30 for *MAPKKK15* and *MAPKKK16*, and 21 for 18S rRNA. (**c**) Germination of FH (DAH 2) and AR seeds (DAH 111-113). The seeds were imbibed for 7 days. The experiments were performed in three independent seed batches with three replicates in each. We obtained similar results from the three experiments, and typical data are presented. The values were presented as mean (±SD) of three technical replicates (n = 8). We could not find significant differences between WT and the mutants (Tukey HSD tests, $P < 0.05$).

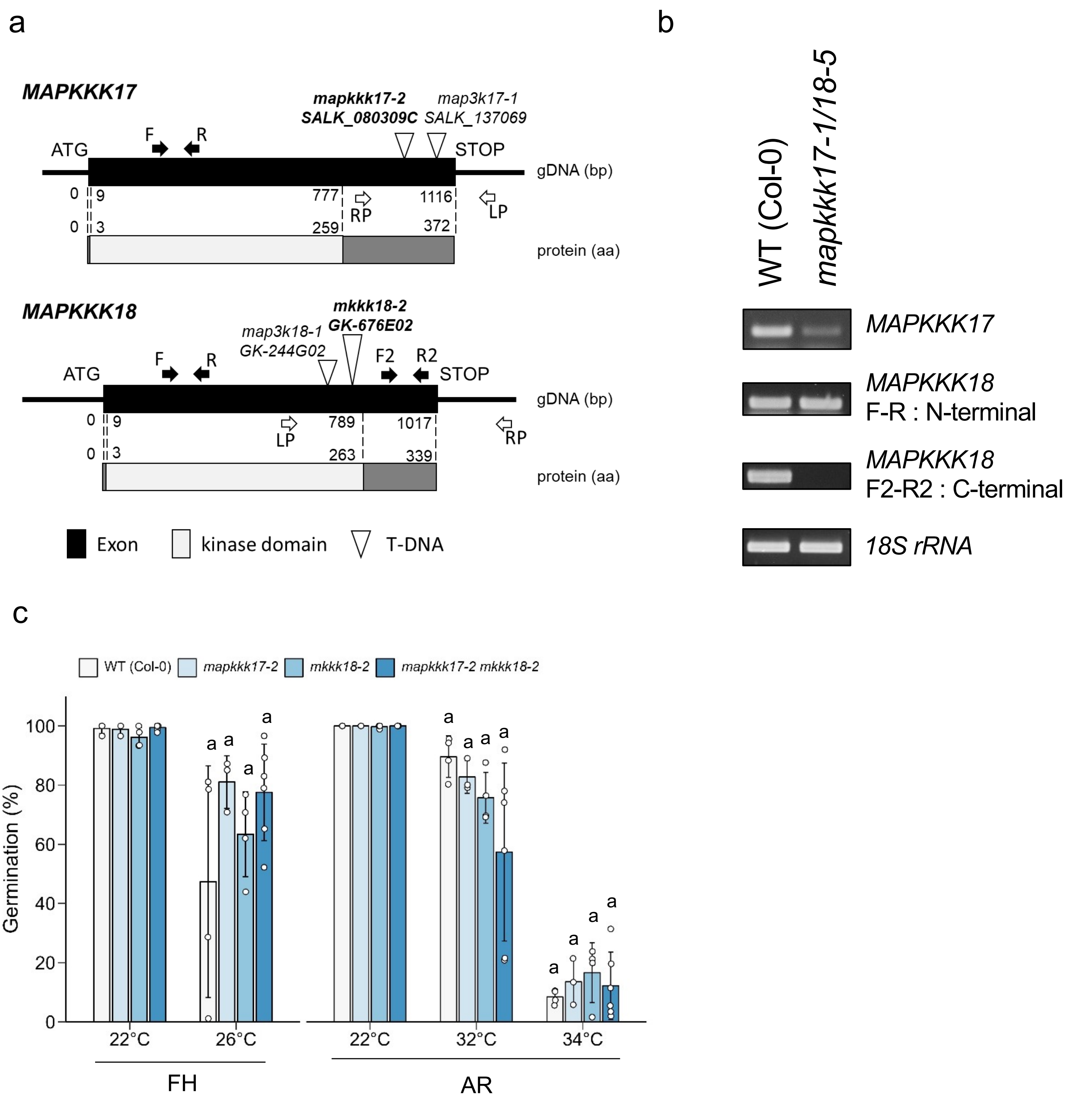


**Supplementary Fig. 8 Germination of loss-of-function mutant seeds of *MAPKKK17* and *MAPKKK18*.** (**a**) Gene model of *MAPKKK17* and *MAPKKK18,* and the positions of T-DNA insertion positions. Positions of primers used for expression analysis are indicated by arrows. (**b**) Semi-quantitative RT-PCR for the mutant gene expression analysis. Total RNA was extracted from 7-day-old seedlings treated with 10 μM ABA for 3 h. 18s rRNA was used as an internal control. PCR cycle numbers were 30 for *MAPKKK17* and *MAPKKK18*, and 21 for 18S rRNA. (**c**) FH (DAH 2) and AR (DAH 80) seeds were imbibed for 7 and 5 days, respectively. We obtained similar results from triplicate experiments, and typical data are presented. We could not find significant differences between WT and the mutants (Tukey HSD tests, P < 0.05).

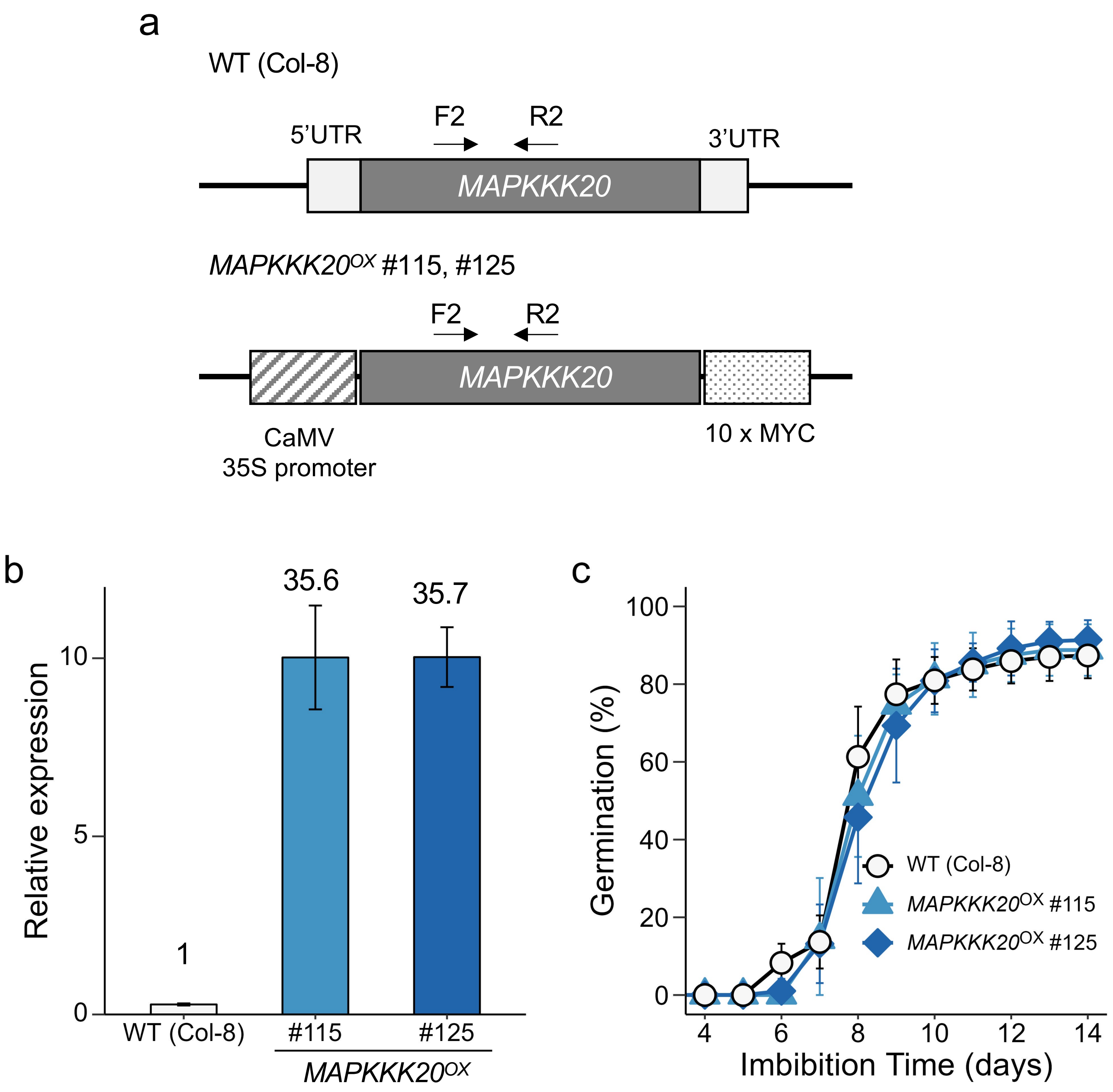


**Supplementary Fig. 9 Over-expression of *MAPKKK20* and the effect on germination at sub-optimal temperature.**
(**a**) Schematic representation of *MAPKKK20*$^{OX}$ construct. Positions of primers used for expression analysis are indicated by arrows. (**b**) Quantification of *MAPKKK20* transcripts by qRT-PCR with At2g20000 as an internal control. Relative expression to WT (Col-8) is indicated by the fold change expression in *MAPKKK20*$^{OX}$ dry seeds. The values are the means (±SD) of three technical replicates. (**c**) Germination time course of AR (DAH 167) seeds imbibed at 5 oC. The values are the mean (±SD) of three biological replicates.

c. Bulk seeds for 6 plants: 30 seeds each x 3 replicates, 3 batch (average of batches)

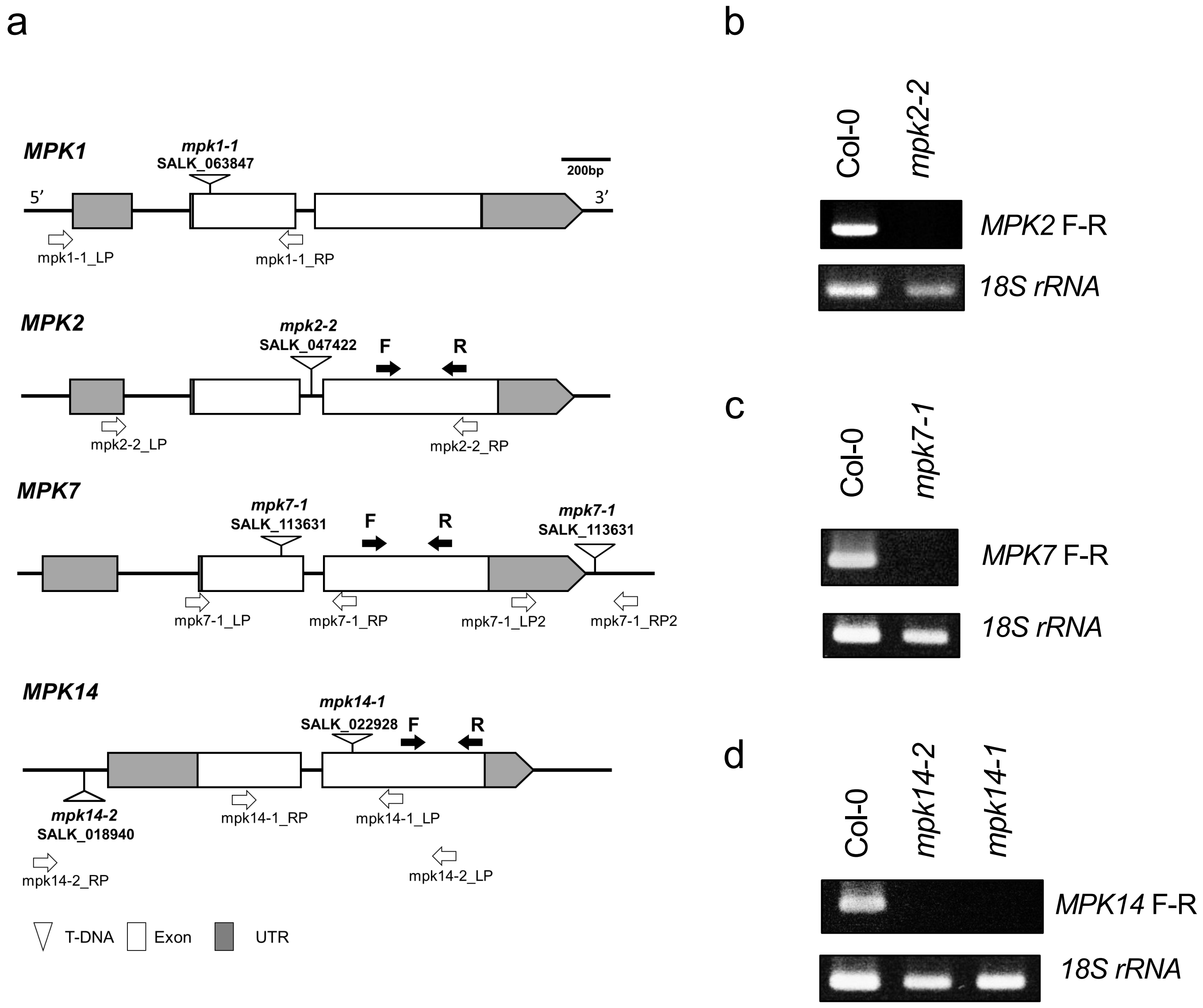


**Supplementary Fig. 10 T-DNA insertion alleles of group C MPKs.**

(**a**) Schematic representation of the genes and position of T-DNA (white triangles) insertion in Col-0. Positions of primers used for genotyping (white arrows) and expression (black arrows) analyses are indicated. (**b-d**) Semi-quantitative RT-PCR for the mutant allele expression analysis. Total RNA was extracted from 24 h imbibed seeds. 18s rRNA was used as an internal control. PCR cycle numbers were 30 for *MPK*s and 21 for 18S rRNA.

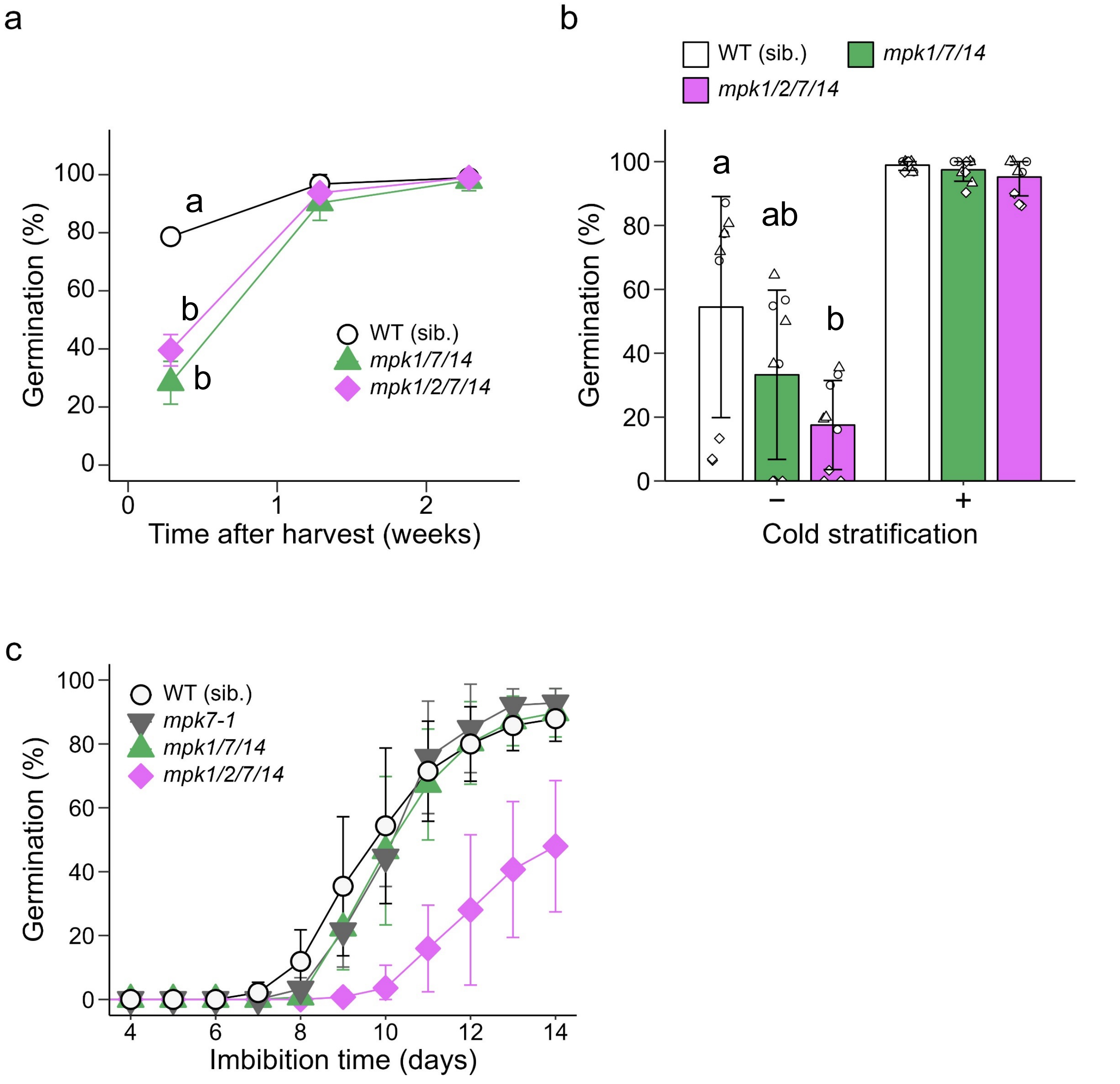


**Supplementary Fig. 11 Seed dormancy and germination of group C MPK multiple mutants at sub-optimal temperature**

(**a**) Enhanced dormancy phenotype of *mpk1/7/14 and mpk1/2/7/14* seeds. Freshly harvested seeds were stored in a desiccator at room temperature. Seeds were imbibed at 22 °C for 7 days without cold stratification. Values are means of three technical replicates with SDs. We obtained similar results from triplicate experiments, and typical data are presented. (**b**) Effect of cold stratification on germination of the freshly harvested seeds. The seeds at DAH 2 were imbibed at 22 °C for 5 days with (+) or without (−) preceding cold stratification at 4 °C for 4 days. (**c**) Germination time course of AR seeds imbibed at 5 °C. (**a, b**) Significant differences between samples are indicated by different letters (Tukey HSD tests, $P < 0.05$). (**b**, **c**) Results from three independent seed batches are shown with averages and SDs.

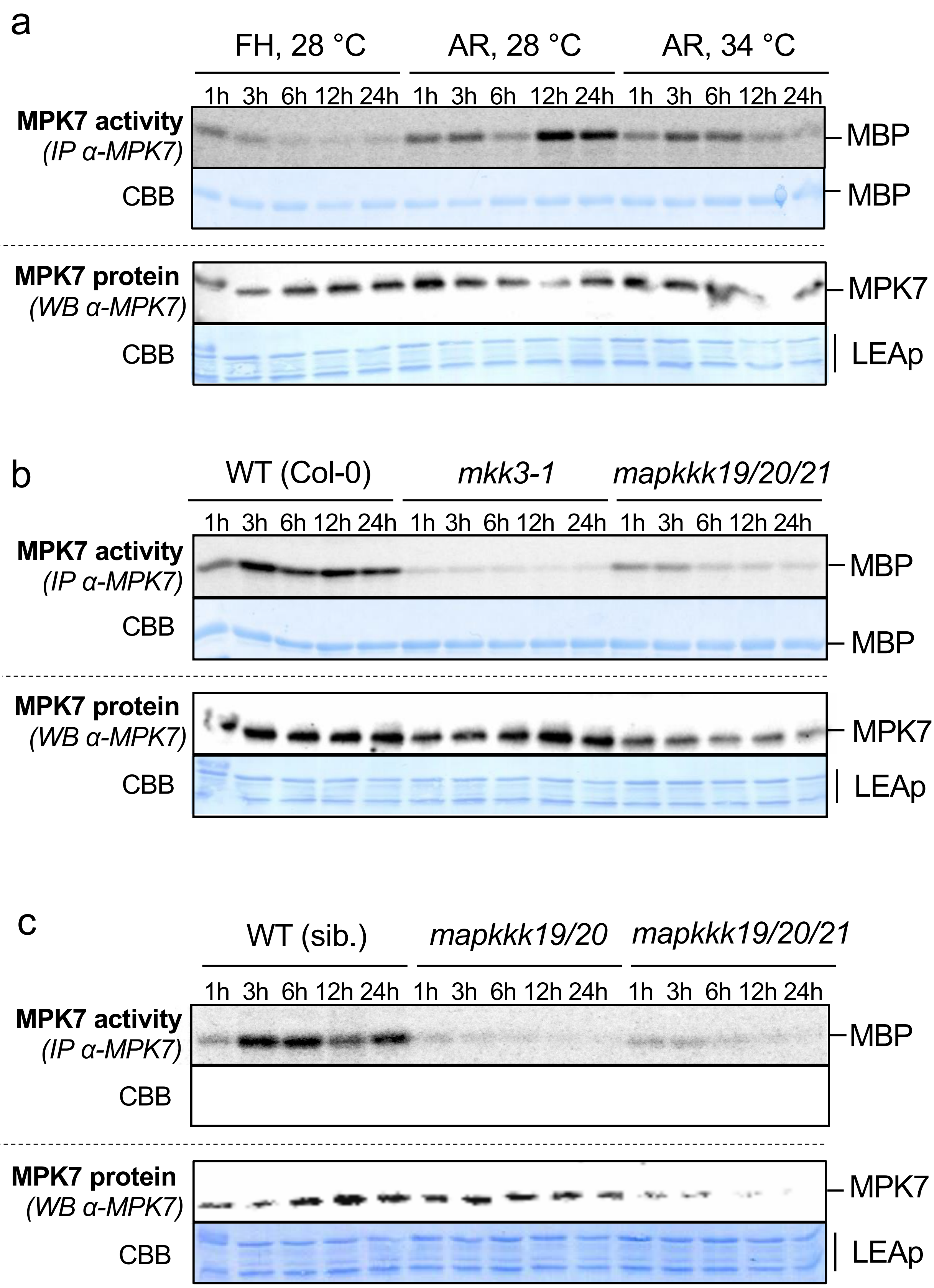


**Supplementary Fig. 12 MPK7 activity in germinating and non-germinating seeds (Biological replication of Fig. 6).** In panel c, electrophoresis and staining of the proteins in the kinase assay mixture has not been done.

**Supplementary Table 1  Genes and its mutant alleles used in this study**

| Allele name | Gene name | AGI code | Stock center code | Stock center | Background | References and notes |
|---|---|---|---|---|---|---|
| *mkk3-1* | *MKK3* | AT5G40440 | SALK_051970 | ABRC, SALK | Col-0 | Takahashi et al. (2007) |
| *mkk3-2* | *MKK3* | AT5G40440 | SALK_208528C | ABRC, SALK | Col-0 | Sözen et al. (2020) |
| *mpk1-1* | *MPK1* | AT1G10210 | SALK_063847C | ABRC, SALK | Col-0 | Enders et al. (2017). Other name: *Atmpk1 (* Ortiz-Masia et al.2007) |
| *mpk2-2* | *MPK2* | AT1G59580 | SALK_047422C | ABRC, SALK | Col-0 | Other name : *mpk2* (Lv et al. 2021) |
| *mpk7-1* | *MPK7* | AT2G18170 | SALK_113631 | ABRC, SALK | Col-0 | This study |
| *mpk14-1* | *MPK14* | AT4G36450 | SALK_022928C | ABRC, SALK | Col-0 | Other name : *mpk14* (Lv et al. 2020) |
| *mapkkk13-1* | *MAPKKK13* | AT1G07150 | GK-277E09 | ABRC, SALK | Col-0 | This study |
| *map3k14-1* | *MAPKKK14* | AT2G30040 | GK-653B01 | ABRC, SALK | Col-0 | gain-of function (Sözen et al. 2020) |
| *mapkkk13/14CR* | *MAPKKK13*<br>*MAPKKK14* | AT1G07150<br>AT2G30040 | -<br>- | | Col-0 | Sözen et al. 2020, Regnard et al. submitted back-to-back |
| *mapkkk15-1* | *MAPKKK15* | AT5G55090 | SALK_084817 | ABRC | Col-0 | Qun et al. 2021 |
| *map3k16* | *MAPKKK16* | AT4G26890 | SALK_003255C | ABRC | Col-0 | Choi et al., 2017 |
| *mapkkk17-2* | *MAPKKK17* | AT2G32510 | SALK_080309C | ABRC | Col-0 | Other name : *mapkkk17* (Romero-Hernandez and Martinez, 2022) |
| *mkkk18-2* | *MAPKKK18* | AT1G05100 | GK-676E02 | ABRC | Col-0 | Mitula et al. 2015 |
| *mapkkk19-1* | *MAPKKK19* | AT5G67080 | pst14411 | RIKEN BRC | Nossen | This study |
| *mapkkk19-1C* | *MAPKKK19* | AT5G67080 | pst14411 | Original | Col-0 | This study. Introgression line of *mapkkk19-1* BC4F2 |
| *mapkkk20-3* | *MAPKKK20* | AT3G50310 | GK-458D07 | ABRC | Col-0 | This study |
| *MAPKKK20* $^{OX}$ | *MAPKKK20* | AT3G50310 | - | | Col-8 | This study<br>[35S::MAPKKK20-10xMyc] |
| *mapkkk21-3* | *MAPKKK21* | AT4G36950 | psh20310 | RIKEN BRC | Nossen | This study |
| *mapkkk21-3C* | *MAPKKK21* | AT4G36950 | psh20310 | Original | Col-0 | Introgression line of *mapkkk21-3* BC3F2 |
| *ga3ox1-3* | *GA3ox1* | AT1G15550 | SALK_004521 | a kind gift from Dr. Yamaguchi | Col-0 | |
| *ga3ox2-1* | *GA3ox2* | AT1G80340 | - | a kind gift from Dr. Yamaguchi | Col-0 | Mitchum et al. (2016) |
| *aba2-2* | *ABA2* | AT1G52340 | - | a kind gift from Dr. Nambara | Col-0 | Nambara et al. (1998) |

**Supplementary Table 2 Primers used in this study**

| Gene name | AGI code | Primer name | Sequence (5'→3') | note |
|---|---|---|---|---|
| Confirmation of T-DNA or transposon insertion | | | | |
| *MAPKKK13* | AT1G07150 | mapkkk13-1_LP | CGGTCCGGTTCCCTCTAT | |
| *MAPKKK13* | AT1G07150 | mapkkk13-1_RP | ATGGTTGGGTGGAAGTTAGATGC | |
| *MAPKKK14* | AT2G30040 | map3k14-1_LP | TGGATCGGCGGTTGAGTTTGA | |
| *MAPKKK14* | AT2G30040 | map3k14-1_RP | CTCGGTTGTCTTACGGAATCATCG | |
| *MAPKKK15* | AT5G55090 | mapkkk15-1_LP | CTTAATTCCCCGTGGAGACTGG | |
| *MAPKKK15* | AT5G55090 | mapkkk15-1_RP | GGAGGTAGCACGTGGTGAAGAA | |
| *MAPKKK16* | AT4G26890 | map3k16_LP | GAGGCTCTACCGCCACTGT | |
| *MAPKKK16* | AT4G26890 | map3k16_RP | AATCCATCCGCCATCGTCTTCC | |
| *MAPKKK17* | AT2G32510 | mapkkk17-2_LP | GATGAATCGTCACATTGGCCTA | |
| *MAPKKK17* | AT2G32510 | mapkkk17-2_RP | GACAGCGACACAACTCCTAAACCA | |
| *MAPKKK18* | AT1G05100 | mapkkk18-5_LP | GAAGCAACGTGTTGGTCGGAG | |
| *MAPKKK18* | AT1G05100 | mapkkk18-5_RP | CCTAACCGAAGCGTACCAAAT | |
| *MAPKKK19* | AT5G67080 | mapkkk19-1_LP | GTACGGTGTTTCGGCGAGGAT | |
| *MAPKKK19* | AT5G67080 | mapkkk19-1_RP | CGATCGTTTGTGAACCGGAAG | |
| *MAPKKK20* | AT3G50310 | mapkkk20-3_LP | CCTTCTTCGGACAACATCTCT | |
| *MAPKKK20* | AT3G50310 | mapkkk20-3_RP | GAAGAAGTTCCACCATAAGGCAC | |
| *MAPKKK21* | AT4G36950 | mapkkk21-3_LP | CGGCGAACCTACCAATGAACC | |
| *MAPKKK21* | AT4G36950 | mapkkk21-3_RP | TTCTCACACCACCCACGTTTCC | |
| *MKK3* | AT5G40440 | mkk3-1_LP | CAAGTCTGATGGTGTTTCCTC | |
| *MKK3* | AT5G40440 | mkk3-1_RP | GAAAGAACCGGCAAGGAAATC | |
| *MKK3* | AT5G40440 | mkk3-2_LP | TTTGAATGCGGCACTGGAGAG | |
| *MKK3* | AT5G40440 | mkk3-2_RP | TGAAATCGGCCAACACGTCTAC | |
| *MPK1* | AT1G10210 | mpk1-1_LP | GATAGAGACGTGAATCTGTTTG | |
| *MPK1* | AT1G10210 | mpk1-1_RP | ATCAACCACACACCTGGAACAAG | |
| *MPK2* | AT1G59580 | mpk2-2_LP | CTCACCAGCCAAATCTCTCCA | |
| *MPK2* | AT1G59580 | mpk2-2_RP | CTCTCTGAACAGTACCTGCAA | |
| *MPK7* | AT2G18170 | mpk7-1_LP | CTTGAGATCGCAGTTCGCATT | |
| *MPK7* | AT2G18170 | mpk7-1_RP | GCGATGTTAGTTGAGCCACCA | |
| *MPK14* | AT4G36450 | mpk14-1_LP | TTGCGTGCGGGTAAATATGAG | |
| *MPK14* | AT4G36450 | mpk14-1_RP | TTTCAGGTACGTTGGTTCGAT | |
| *GA3OX1* | AT1G15550 | ga3ox1-3_LP | GATACTCTTTCCATGTCACCGA | |
| *GA3OX1* | AT1G15550 | ga3ox1-3_RP | CATTCACCTCCCACACTCTC | |
| *GA3OX2* | AT1G80340 | ga3ox2-1_LP | AGATCATTATATCGGATGGTGG | |
| *GA3OX2* | AT1G80340 | ga3ox2-1_RP | GCCTTTTAGCATGAGTTCAACG | |
| T-DNA | - | LB_6313R for SALK | TCAAACAGGATTTTCGCCTGCT | SALK T-DNA pROK2 left border primer |
| T-DNA | - | GABI_LB | CATATTGACCATCATACTCATTGCTG | GABI T-DNA pAC161 left border primer |
| T-DNA | - | LB_pNU74 | ACTGGCATGACGTGGGTTTCT | pNU74 left border primer used for genotyping of *ga3ox2-1* |
| Transposon | - | Ds3-2a | CCGGATCGTATCGGTTTTCG | RIKEN BRC Ds-transposon H-edge primer |
| Vector | - | pGWB20_qRT-PCR_R | GTTCACCGTTAATTAACCCGCTG | used for genotyping of MAPKKK20OX |
| qRT-PCR | | | | |
| *MAPKKK19* | AT5G67080 | MAPKKK19_qRT-PCR_F | CAGAGGACGTCTCGACATCGC | |
| *MAPKKK19* | AT5G67080 | MAPKKK19_qRT-PCR_R | ACCGTACGGTGACCCAGCTATTG | |
| *MAPKKK20* | AT3G50310 | MAPKKK20_qRT-PCR_F | TGGTCTGTCGGTGGGAGTTG | |
| *MAPKKK20* | AT3G50310 | MAPKKK20_qRT-PCR_R | GTAGAATGGGTCTAACTAGGTGCT | |
| *MAPKKK20* | AT3G50310 | MAPKKK20_qRT-PCR_F2 | ACGCTAAAGGGTTTGCACACTG | also used for genotyping of MAPKKK20OX |
| *MAPKKK20* | AT3G50310 | MAPKKK20_qRT-PCR_R2 | CATGTATAACGGCGTCCCTCTAA | |
| *MAPKKK21* | AT4G36950 | MAPKKK21_qRT-PCR_F | ACACGACACCGACCAATCAGG | |
| *MAPKKK21* | AT4G36950 | MAPKKK21_qRT-PCR_R | ATTGGTAGGTTCGCCGGAGGA | |
| *NCED2* | AT4G18350 | NCED2_qRT-PCR_F | GCGGCTGAGCGTGCATTAA | |
| *NCED2* | AT4G18350 | NCED2_qRT-PCR_R | GGGAATAATTCCCGGCAATCT | |
| *NCED5* | AT1G30100 | NCED5_qRT-PCR_F | CCTCCGTTAGTTTCACCAACACT | |
| *NCED5* | AT1G30100 | NCED5_qRT-PCR_R | GGTGTGTCGGAGACGGAGTT | |
| *NCED9* | AT1G78390 | NCED9_qRT-PCR_F | GGAAAACGCCATGATCTCACA | |
| *NCED9* | AT1G78390 | NCED9_qRT-PCR_R | AGGATCCGCCGTTTTAGGAT | |
| *GA3ox1* | AT1G15550 | GA3ox1_qRT-PCR_F | GGACAAACCGGGTAGTGATTT | |
| *GA3ox1* | AT1G15550 | GA3ox1_qRT-PCR_R | ATGGGCCAGTCTCAGTTCAG | |
| *GA3ox2* | AT1G80340 | GA3ox2_qRT-PCR_F | GTTTCCAAGTGAGAGATGGGTAG | |
| *GA3ox2* | AT1G80340 | GA3ox2_qRT-PCR_R | TCTCCACTTCCCAAACTGGT | |
| *CYP707A1* | AT4G19230 | CYP707A1_qRT-PCR_F | CTCACTCTCTTCGCCGGAAG | |
| *CYP707A1* | AT4G19230 | CYP707A1_qRT-PCR_R | GGAGGGAGTGGGAGTTTGGAA | |
| *CYP707A2* | AT2G29090 | CYP707A2_qRT-PCR_F | CGTCTCTCACATCGAGCTCCTT | |
| *CYP707A2* | AT2G29090 | CYP707A2_qRT-PCR_R | GAGGGTGTTGATGGACTTTTGG | |
| *CYP707A3* | AT5G45340 | CYP707A3_qRT-PCR_F | CTCTGTTTCTCTGTTTACTCCGATTTA | |
| *CYP707A3* | AT5G45340 | CYP707A3_qRT-PCR_R | CGTATCTTCTCTGTTTTGCTGCA | |
| *HBT* | AT2G20000 | At2g20000_qF1 | GTATAGCTCCACCACCACTT | reference genes for transcript normalization |
| *HBT* | AT2G20000 | At2g20000_qR1 | TCTTCTAGGTGCTTGAAGAGT | reference genes for transcript normalization |
| semi-qRT-PCR | | | | |
| *MAPKKK13* | AT1G07150 | MAPKKK13_F | GTGCAGTTAGTACGGCGATCAG | |
| *MAPKKK13* | AT1G07150 | MAPKKK13_R | AGGCTTTAGCGAGCGGAAGACA | |
| *MAPKKK15* | AT5G55090 | MAPKKK15_F1 | CTTACATAGTCAAGTACATTGGTTC | N-terminal |
| *MAPKKK15* | AT5G55090 | MAPKKK15_R2 | CCGATCATCACATTCTGGCTCT | N-terminal |
| *MAPKKK15* | AT5G55090 | MAPKKK15_F2 | ACGAACGGTTGGATTGAAGTAAG | C-terminal |
| *MAPKKK15* | AT5G55090 | MAPKKK15_R2 | AAATAATAAATGTTGTCCTCAGGGG | C-terminal |
| *MAPKKK16* | AT4G26890 | MAPKKK16_F | TAACGCGCGAGAGCAACGGA | |
| *MAPKKK16* | AT4G26890 | MAPKKK16_R | CTTCAAGTCGCAGTGAACAATCC | |
| *MAPKKK17* | AT2G32510 | MAPKKK17_F | AGGTTCACCGCTCAGAGTTC | |
| *MAPKKK17* | AT2G32510 | MAPKKK17_R | CATACGGTGCGTACTCCATC | |
| *MAPKKK18* | AT1G05100 | MAPKKK18_F | CTCCGTACGTCATCGGATACA | N-terminal |
| *MAPKKK18* | AT1G05100 | MAPKKK18_R | AAGCCGCCGTTCTTGGTCG | N-terminal |
| *MAPKKK18* | AT1G05100 | MAPKKK18_F2 | AGCGAGTCAACTACTAAACCATC | C-terminal |
| *MAPKKK18* | AT1G05100 | MAPKKK18_R2 | GACTAATTCCGTCGAACCGTG | C-terminal |
| *MAPKKK19* | AT5G67080 | MAPKKK19_F | CAGAGGACGTCTCGACATCGC | |
| *MAPKKK19* | AT5G67080 | MAPKKK19_R | ACCGTACGGTGACCCAGCTATTG | |
| *MAPKKK20* | AT3G50310 | MAPKKK20_F | TGGTCTGTCGGTGGGAGTTG | |
| *MAPKKK20* | AT3G50310 | MAPKKK20_R | GTAGAATGGGTCTAACTAGGTGCT | |
| *MAPKKK21* | AT4G36950 | MAPKKK21_F1 | ACACGACACCGACCAATCAGG | C-terminal |
| *MAPKKK21* | AT4G36950 | MAPKKK21_R1 | ATTGGTAGGTTCGCCGGAGGA | C-terminal |
| *MAPKKK21* | AT4G36950 | MAPKKK21_F2 | CTAATGGAGTGGATTCGTAGAG | N-terminal |
| *MAPKKK21* | AT4G36950 | MAPKKK21_R2 | CCGAACAGTCACCGAGATCA | N-terminal |
| *MPK2* | AT1G59580 | MPK2-F1 | GTACCGAGCACCAGAGCTAC | |
| *MPK2* | AT1G59580 | MPK2-R1 | AAGCGCTTCCGTGACACTAATC | |
| *MPK7* | AT2G18170 | MPK7-F | CATGTGCAGGCGGATTGGAT | |
| *MPK7* | AT2G18170 | MPK7-R | GGTTGGTTGCATATTCGCGG | |
| *MPK14* | AT4G36450 | MPK14-F | TTAAGCTCGGGGGAGGTAATG | |
| *MPK14* | AT4G36450 | MPK14-R | CTCATATTTACCCGCACGCAA | |
| *18SrRNA* | AT3G41768 | rt-18S-F | ATACGTGCAACAAACCC | reference gene |
| *18SrRNA* | AT3G41768 | rt-18S-R | CTACCTCCCCGTGTCA | reference gene |
| Cloning of MAPKKK20 | | | | |
| *MAPKKK20* | AT3G50310 | OMN126_MAPKKK20_attb_Fwd | GGGGACAAGTTTGTACAAAAAAGCAGGCTCCATGGAGTGGGTTCGAGGAGAAA | Gateway MAPKKK20 Fwd primer |
| *MAPKKK20* | AT3G50310 | OMN127_MAPKKK20_attb_Rev | GGGGACCACTTTGTACAAGAAAGCTGGGTCCCGGACTGTAAGCCAACTC | Gateway MAPKKK20 Rev primer |
